# Whole-brain dynamics of human sensorimotor adaptation

**DOI:** 10.1101/2020.11.27.401679

**Authors:** Dominic I. Standage, Corson N. Areshenkoff, Daniel J. Gale, Joseph Y. Nashed, J. Randall Flanagan, Jason P. Gallivan

## Abstract

Humans vary greatly in their motor learning abilities, yet little is known about the neural processes that underlie this variability. We identified distinct profiles of human sensorimotor adaptation that emerged across two days of learning, linking these profiles to the dynamics of whole-brain functional networks early on the first day, when cognitive strategies toward sensorimotor adaptation are believed to be most prominent. During early learning, greater recruitment of a network of higher-order brain regions, involving prefrontal and anterior temporal cortex, was associated with faster learning. At the same time, greater integration of this ‘cognitive network’ with a sensorimotor network was associated with slower learning, consistent with the notion that cognitive strategies toward adaptation operate in parallel with implicit learning processes of the sensorimotor system. On the second day, greater recruitment of a network that included the hippocampus was associated with faster re-learning, consistent with the notion that savings involves declarative memory systems. Together, these findings provide novel evidence for the role of higher-order brain systems in driving individual differences in adaptation.

## Introduction

Successful behaviour relies on the brain’s ability to flexibly adapt to environmental changes by learning new sensorimotor mappings. For example, a tennis player may favour a particular style of racket, but she will quickly adjust to a racket with a longer handle. For decades, it was widely believed that such sensorimotor adaptation was largely implicit, driven by sensory prediction errors reflecting the difference between the expected and sensed consequences of movement (Wolpert and Ghahramani 2000). Recent studies, however, have shown that explicit (declarative) cognitive strategies can also play an important role in guiding these adjustments, operating in parallel with implicit learning (see Krakauer and Mazzoni 2011; McDougle, Ivry, and Taylor 2016). These two learning components have been well characterised behaviourally and have been shown to operate on different timescales, with slow and fast adaptation linked to the implicit and explicit components respectively (Smith, Ghazizadeh, and Shadmehr 2006; McDougle, Bond, and Taylor 2015). There is emerging behavioural evidence that individual differences in the initial rate of adaptation are driven by the explicit component (Fernandez-Ruiz et al. 2011; de Brouwer et al. 2018), with the implicit component exhibiting minimal variability across individuals. Moreover, it has been suggested that the explicit component is responsible for the phenomenon of savings, or faster re-learning upon reexposure to the task (Morehead et al. 2015; Haith, Huberdeau, and Krakauer 2015). While a large body of work indicates that the cerebellum is essential for implicit adaptation (Martin et al. 1996; Tseng et al. 2007; Izawa, Criscimagna-Hemminger, and Shadmehr 2012), the neural bases of explicit adaptation remain poorly understood (McDougle, Ivry, and Taylor 2016).

Several previous studies have used functional neuroimaging methods to document changes in the activity of brain regions during sensorimotor adaptation (R. Shadmehr 1997; Anguera et al. 2007; Doyon et al. 2009; Dayan and Cohen 2011; Krakauer et al. 2004), including time-resolved changes (Bédard and Sanes 2014). It is widely acknowledged, however, that this type of learning is supported by distributed brain circuitry, spanning the cerebrum, cerebellum and striatum (Doyon et al. 2009). Understanding such distributed neural systems requires an investigation of the interactions between brain regions in large-scale networks (R. Shadmehr 1997; Anguera et al. 2007; Doyon et al. 2009; Dayan and Cohen 2011; Krakauer et al. 2004).

In recent years, the use of graphical methods to construct and analyse networks derived from functional magnetic resonance imaging (fMRI) data has been an informative approach for studying large-scale brain networks (Sporns 2014; Bullmore and Sporns 2009a). Under this *network neuroscience* approach (Bassett and Sporns 2017), coactive brain regions are hypothesised to form a functional network during a time window of interest. A compelling aspect of these networks is their modularity, or their decomposition (a.k.a. partitioning) into subnetworks, characterised by denser (sparser) connectivity between regions in the same (different) module(s). Modularity has long been recognised as a crucial property of complex biological systems (see Bassett et al. 2011) and the structural and functional organisation of the human brain is highly modular (Bullmore and Sporns 2009b; Ferrarini et al. 2009). An important development in this field — crucial for examining dynamic processes like learning — is the construction and analysis of temporally dynamic functional brain networks (Bassett and Sporns 2017). For instance, recent fMRI studies have shown that the propensity of participants’ brain regions to change modular affiliation over time (known as flexibility) correlates with cognitive task performance (Braun et al. 2015) and motor skill acquisition (Bassett et al. 2011).

We used this dynamic network neuroscience approach to investigate the whole-brain bases of sensorimotor adaptation. We collected fMRI data while participants performed a classic sensorimotor adaptation task over two days, separated by twenty-four hours. We measured participants’ whole-brain modular reconfiguration, as well as the recruitment (construction) and integration of functional networks derived from the dynamic modules, asking whether patterns of learning over both days of the experiment could be predicted by these network measures during early learning on the first day, when cognitive strategies during sensorimotor adaptation are believed to be most prominent (de Brouwer et al. 2018; Taylor, Krakauer, and Ivry 2014).

## Results

### Overview of approach

On two testing days, separated by 24 hours, we recorded whole-brain fMRI activity while participants (N=40; 32 used in our analysis) adapted, de-adapted, and then re-adapted (on the second day) their movements during a visuomotor rotation (VMR) task (Krakauer 2009). This error-based learning task required participants to use their right (dominant) hand to perform centre-out movements of a cursor to single targets, pseudo-randomly selected without replacement from 8 possible locations. To establish baseline performance, participants executed 120 standard, non-rotation trials (i.e. 15 bins of 8 trials). Following this baseline condition (henceforth baseline), a 45° clockwise rotation was applied for the next 320 trials (henceforth learning), where the viewed cursor was rotated about the hand’s start location, requiring that participants learn to adapt their movements to the perturbation (see Fig. 1A). Finally, to allow for an investigation of relearning the following day, we de-adapted participants’ learning with 120 more baseline trials (henceforth washout) (Krakauer 2009). One day later, participants returned to undergo the same testing regimen, enabling us to measure savings (i.e. faster re-learning on Day 2 than Day 1) upon re-exposure to the VMR (Morehead et al. 2015). At the behavioural level, this approach enabled us to examine participants’ initial rates of learning (Day 1) and re-learning (Day 2), as well as their expression of savings across days. At the neural level, this approach enabled us to examine the dynamics of whole-brain modules during learning, as well as the regional composition of these modules.

**Figure 1.**
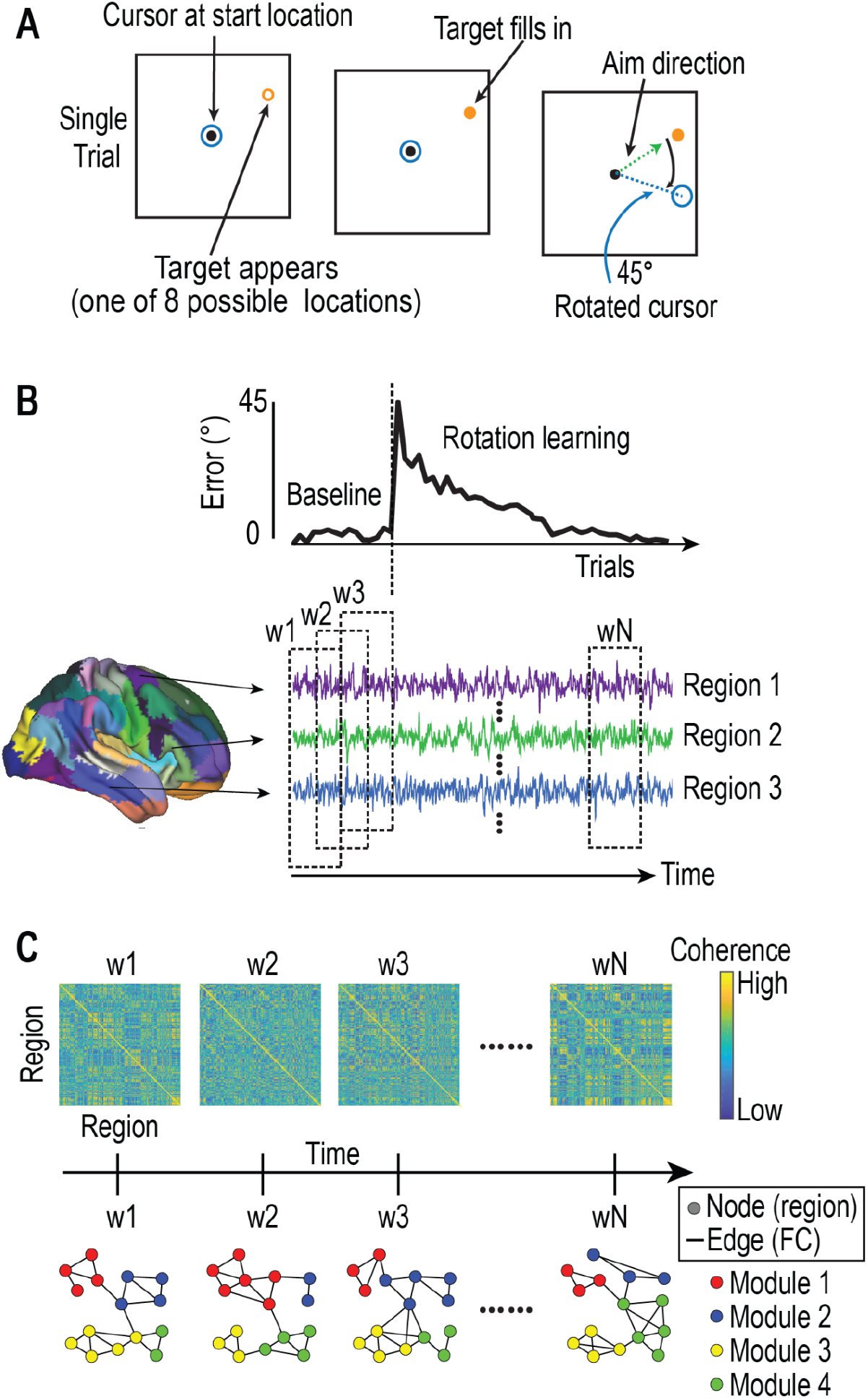
Overview of task and neural analyses. (A) Visuomotor rotation (VMR) task, where the viewed cursor, controlled by the hand, is rotated about the movement start location. (B) Each participant’s cerebrum, striatum and cerebellum were parcellated into discrete brain regions and the average % Blood Oxygenation Level-Dependent (BOLD) time series was extracted from each region for the entire task on each day (three example cerebral regions shown). The coherence (of scale 2 wavelet coefficients, see Methods) was calculated for each pair of regions in sliding, half-overlapping windows, constructing whole-brain functional connectivity matrices for each window (w1-wN; shown in ‘C’). (C) Time-resolved clustering methods were applied to the functional connectivity matrices constructed in ‘B’, linking module labels (for each node) across time slices. This procedure allowed us to identify network modules that evolved during learning (4 modules shown in the schematic).

### Behavioural analyses reveal three distinct profiles of learning across days

Group-level behaviour was consistent with earlier studies, where (on average) participants learned to reduce their errors during task performance and exhibited savings (Krakauer 2009; Morehead et al. 2015). The left panel in Figure 2A shows mean learning curves and savings over all participants. These group-level findings, however, obscure significant inter-participant variability in the patterns of learning across days. For example, participant P1 (Figure 2B) showed fast learning on both days (fast-fast, FF), participant P2 showed slow learning on both days (slow-slow, SS) and participant P3 showed a pattern of slow learning on Day 1 (magenta trace) but fast learning on Day 2 (cyan trace, i.e., slow-fast, SF). To determine whether these very different patterns of learning were extreme examples from a spectrum or reflected a limited number of distinct learning profiles, we performed *k*-means clustering across all participants’ early and late binned errors on each day, along with their savings (*i*.*e*. five behavioural measures per participant). These measures captured each participant’s rate of learning (early binned error on each day), completeness of learning (late binned error on each day) and improvement in learning rate between days (savings) respectively. The clustering algorithm separated participants into three distinct subgroups or *profiles* of learners (Figure 2C) that showed very similar patterns of learning (Figure 2D) to the FF, SS and SF participants shown in Figure 2B (15 participants were FF, 10 were SS and 7 were SF). Our clustering solution of the subgroups was statistically significant according to the silhouette (p=0.004) and Calinksi-Harabasz (p=0.009) indices, the same two indices used to determine the optimal number of clusters (Figure 2C). Its cluster distance matrix is shown in Figure 2G and the procedure for determining statistical significance is described in Methods.

**Figure 2.**
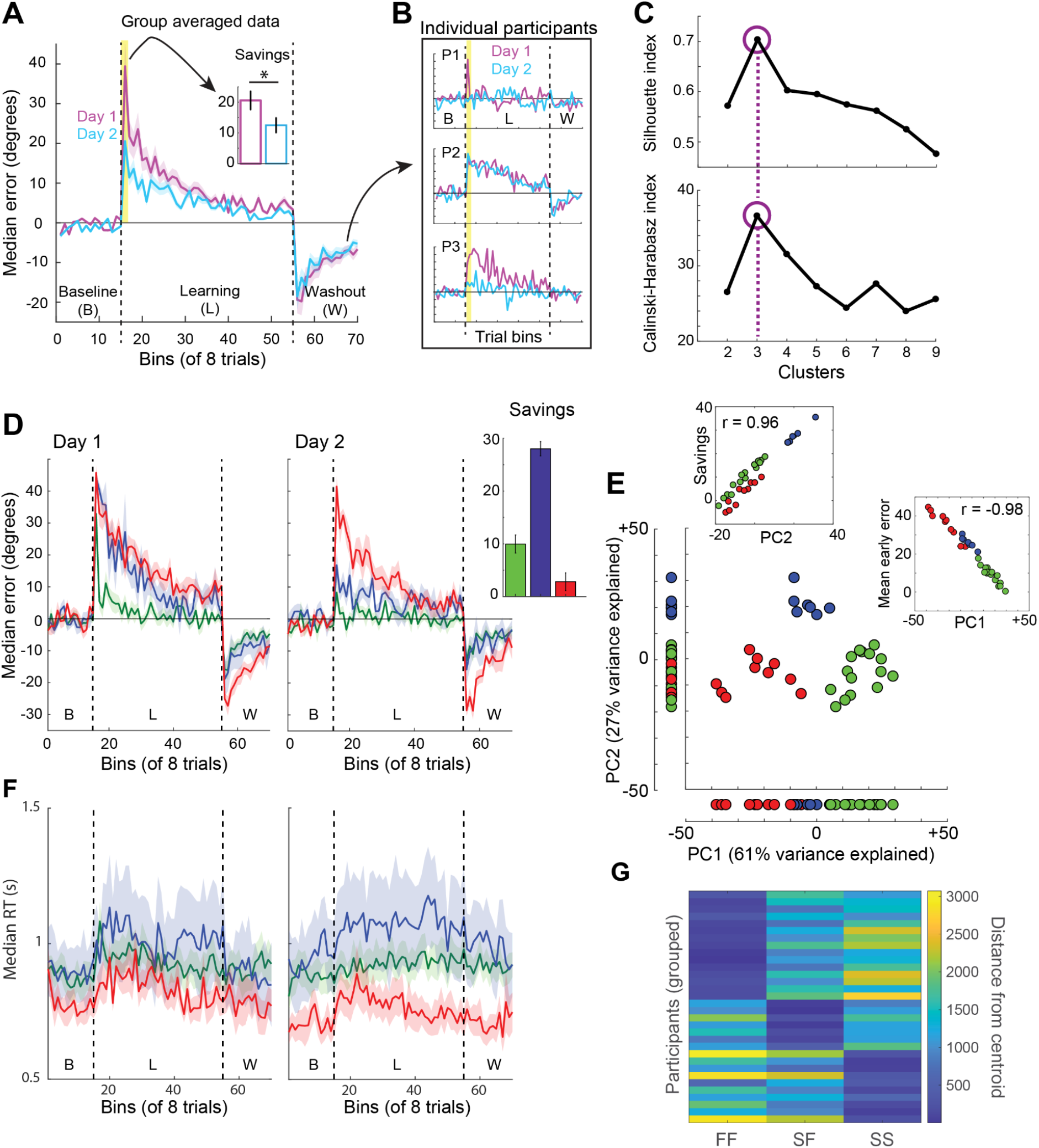
Clustering of participants’ learning data revealed 3 behavioural subgroups. **(A)** Mean (across participants) of the binned median error (in degrees) during baseline (non-rotation), learning (45 degree rotation of the cursor) and washout (non-rotation) on Day 1 (magenta) and Day 2 (blue). Shading shows standard error (±1 SE) and the dashed vertical lines demark the three task components. Note that savings (inset) is significant at the group level [paired t-test; t(31)=6.122, p=8.666e-7]. **(B)** The group-averaged approach in panel A obscures differences in learning between +_individuals (3 example participants, P1-P3) **(C)** *k*-means clustering of participants’ early and late errors and savings across Days 1 and 2 (5 variables in total) identified three subgroups of participants. Plot shows the silhouette and Calinski-Harabasz (C-H) indices (see Methods), measuring the goodness of clustering. Both measures are maximized by three clusters (purple circles and dashed line indicate the best clustering solution). **(D)** The mean (across participants) binned median error for the three behavioural subgroups (identified in B,C) during baseline, learning and washout on Day 1 (left) and Day 2 (right). The subgroups are readily distinguished by their early error on each day and we refer to their patterns of learning as fast-fast (FF, green, N=15), slow-slow (SS, red, N=10) and slow-fast (SF, blue, N=7). On Day 1, mean early error of FF was significantly lower than that of SS [two-sample t-test, t(23)=-9.963, p=8.203e-10] and SF [t(20)=-12.74, p=4.691e-11], but there was no statistical difference between SF and SS [t(15)=1.0931, p=0.292]. On Day 2, mean early error of FF was significantly lower than that of SS [t(23)=-7.413, p=8.476e-10] and SF [t(20)=-3.106, p=0.006], and mean early error of SF was significantly lower than that of SS [t(15)=-6.67, p=7.495e-6]. Inset shows savings by behavioural group. The FF and SF subgroups showed significant savings [t-test versus zero; FF: t(14)=5.817, p=4.473e-5; SF: t(6)=20.703, p=8.265e-7], but SS did not [t-test versus zero: t(9)=1.688, p=0.126]. **(E)** Top two components of the PCA for the 5 learning measures across participants. Single data points correspond to participants, colour-coded by their cluster-assigned subgroup. The x-axis of the plot shows that PC1 accurately classifies 31 of 32 participants (97% accuracy). Inset scatter plots show that PC1 and PC2 have strong linear relationships with mean early error across days and savings respectively. **(F)** Mean bin median reaction time (RT) for each subgroup for the baseline, learning and washout epochs on Day 1 (left) and Day 2 (right). Shading shows standard error (±1 SE). **(G)** Cluster distance matrix, showing Euclidian distance from cluster centroids for each participant.

Further examination of these subgroups (Figure 2D) revealed two features of across-day learning that were obscured at the group level. Firstly, learning on Day 1 did not always predict learning on Day 2, since a sizable proportion of ‘slow’ learners on Day 1 (7 of 17 participants, ∼40%) became ‘fast’ learners on Day 2 (*i*.*e*. the SF subgroup). Secondly, savings was not ubiquitous across participants, as suggested by the group-level data in Figure 2A. Rather, savings differed significantly between the subgroups (see Figure 2D inset), with the largest magnitude of savings (∼28°) expressed by SF participants, followed by FF participants (∼10°). In contrast, the SS subgroup did not show significant savings (Figure 2D and caption). Together, these results indicate that a large portion of the savings observed at the group level was driven by the SF subgroup, since FF participants were already fast learners on Day 1 and therefore had limited scope for improvement.

Another unexpected feature of our clustering analysis was that SF participants showed a very similar rate of *de-adaptation* (during washout) to FF participants on Day 1 (Figure 2D), despite having shown a very similar rate of adaptation to SS participants. One interpretation of this finding is that SF participants identified the appropriate strategy for the task (re-aiming in a direction that counteracted the cursor rotation) at some point during Day 1 and could therefore more readily discard that strategy once the rotation was removed (at least at a similar rate to FF participants). This interpretation also helps to explain the large savings exhibited by SF participants, given that savings in visuomotor rotation tasks is thought to reflect the recall of an effective re-aiming strategy (Morehead et al. 2015). Notably, prior work has also provided evidence that strategic re-aiming during sensorimotor adaptation entails longer reaction times (RTs) (Fernandez-Ruiz et al. 2011; Haith, Huberdeau, and Krakauer 2015; de Brouwer et al. 2018). RTs for each behavioural subgroup are shown in Figure 2F, where the means were highest in the SF subgroup and lowest in the SS subgroup. Indeed, we re-ran the above analysis with RT on each day included in the clustering (in addition to the five error-based measures described above) and found that the behavioural subgroups identified under this approach were identical to those described above. This convergence of observations suggests that SF participants were using cognitive strategies on Day 1 (albeit unsuccessfully, see also Miyamoto, Wang, and Smith 2020), despite showing much slower adaptation than FF participants. If so, we reasoned that SF and FF participants may exhibit similar neural data on Day 1, despite their stark differences in task performance. In other words, although SF participants’ learning performance was indistinguishable from that of SS participants on Day 1, SF may exhibit different brain dynamics than SS, foreshadowing their Day 2 task performance.

Next, having defined the three behavioural subgroups (FF, SS and SF), we sought to identify a single scalar measure that accurately captured this subgroup membership, thus avoiding issues of statistical power related to the subgroup sizes and enabling the examination of brain-behaviour relationships with standard correlation methods. To this end, we performed principal component analysis (PCA) on the same five behavioural measures used to derive the subgroups in our clustering analysis above. The first principal component (PC1) accounted for 61% of the variance in the behavioural data and accurately classified 31/32 participants’ subgroup membership (Figure 2E), making it a suitable proxy measure for the subgroups. Guided by the PCA factor loadings (Figure S1), we found that PC1 corresponds to participants’ mean early error across days (r = −0.98, Figure 2E inset), providing an intuitive interpretation of its meaning in relation to learning performance and subgroup membership. We reversed the sign of PC1, so that higher scores correspond to faster learning (and smaller errors). Of note, we found that the second principal component (PC2) closely corresponds to savings (r = 0.96, Figure 2E inset) and accounts for 27% of the variance in the data. While PC2 does not feature in our subsequent analyses, we wish to emphasise that the scatter plot of the first two principal components in Figure 2E demonstrates that the subgroups correspond to highly distinct clusters in a space that accounts for 88% of the total variance in the data, where the axes of this space correspond to interpretable behavioural measures.

With a view to our analysis of participants’ neural data, the above behavioural findings generate two competing hypotheses. If Day 1 neural data effectively indexes participants’ learning performance, then the whole-brain dynamics of SS and SF participants should be similar, differing from FF participants. In contrast, if both the FF and SF subgroups were using cognitive strategies during Day 1 learning — the latter group unsuccessfully (Miyamoto, Wang, and Smith 2020) — then the two groups may nevertheless exhibit similar brain dynamics, despite the stark difference in their Day 1 performance.

### Coordinated modular reconfiguration is associated with faster learning

We constructed multislice networks from each participant’s fMRI data during baseline and learning on each day (see Methods). In each of these task epochs, we partitioned the networks into spatio-temporal modules that maximised a quality function Q (Mucha et al. 2010) and verified that participants’ whole-brain networks showed significant modularity (Bassett et al. 2011, 2015) (see Supplementary Materials). In other words, participants’ whole-brain functional networks were composed of interacting subnetworks. Next, we divided learning into early and late epochs on each day (windows 1 to 14 and 26 to 39 respectively, corresponding to the first and last 14 windows of learning, equal to the 14 baseline windows) and quantified three measures of modular reconfiguration, known as flexibility, cohesion strength (a.k.a. cohesive flexibility) and disjointedness (a.k.a. disjointed flexibility) (Telesford et al. 2017). Flexibility is a general measure that captures the degree to which brain regions change their modular affiliation over time, and has previously been associated with cognitive performance (Braun et al. 2015). Where flexibility captures *that* brain regions change modular affiliation, cohesion strength and disjointedness characterise *how* they change affiliation. More specifically, cohesion strength captures the degree to which brain regions change modules in coordinated groups, and has previously been associated with motor skill acquisition (Telesford et al. 2017). Conversely, disjointedness captures the degree to which modular reconfiguration is *un*coordinated, *i*.*e*. the degree to which brain regions change affiliation on their own, and has previously been associated with unconsciousness (Standage et al. 2020). These measures enabled us to characterise modular reconfiguration during early learning on Day 1, and to determine its relationship with PC1, our proxy for subgroup membership.

During early learning on Day 1 (henceforth early learning) we did not find a significant correlation between PC1 and flexibility (r = −0.015, p = 0.935), indicating that mere changes in modular composition were unrelated to participants’ learning profiles at this time. To determine whether *coordinated* modular changes were related to learning, we calculated the correlation between PC1 and cohesion strength during early learning, finding a significant positive correlation (Figure 3A). We reasoned that if these coordinated changes were directly related to the use of a cognitive strategy, then the correlation would be stronger during early learning than during other task epochs, *i*.*e*. it would be stronger when cognitive strategies are most pronounced (Taylor et al. 2014; McDougle et al. 2015). We therefore used Williams’ test (Steiger 1980) to compare it to the correlation between PC1 and cohesion strength observed during baseline on Day 1, late learning on Day 1 (henceforth late learning), baseline on Day 2, early learning on Day 2 (henceforth early re-learning) and late learning on Day 2 (henceforth late re-learning). Following FDR correction, the correlation was not significantly stronger than in any other epoch [1-tailed Williams test, Day 1 baseline: t(29) = 1.622, p = 0.058, adjusted p = 0.109; late learning: t(29) = 2.368, p = 0.012, adjusted p = 0.062; Day 2 baseline: t(29) = 1.392, p = 0.087, adjusted p = 0.109; early re-learning: t(29) = 1.261, p = 0.109, adjusted p = 0.109; late re-learning: t(29) = 1.327, p = 0.097, adjusted p = 0.109] suggesting that coordinated modular reconfiguration is not directly related to the explicit component (*e*.*g*. the search for a strategy). Rather, it may reflect a more general cognitive property (*e*.*g*. attention) that in turn facilitates fast learning (*e*.*g*. Yerkes and Dodson 1908). Regardless, this property appears to be a global one, as 133 out of the 142 regions in our study showed a positive correlation between PC1 and cohesion strength, 32 of which (22.5%) were significant (0.351 < r < 0.626, 1.294e-4 < p < 0.049). In contrast, none of the remaining 9 negative correlations with PC1 were significant (−0.167 < r < −0.002, 0.359 < p < 0.993).

**Figure 3.**
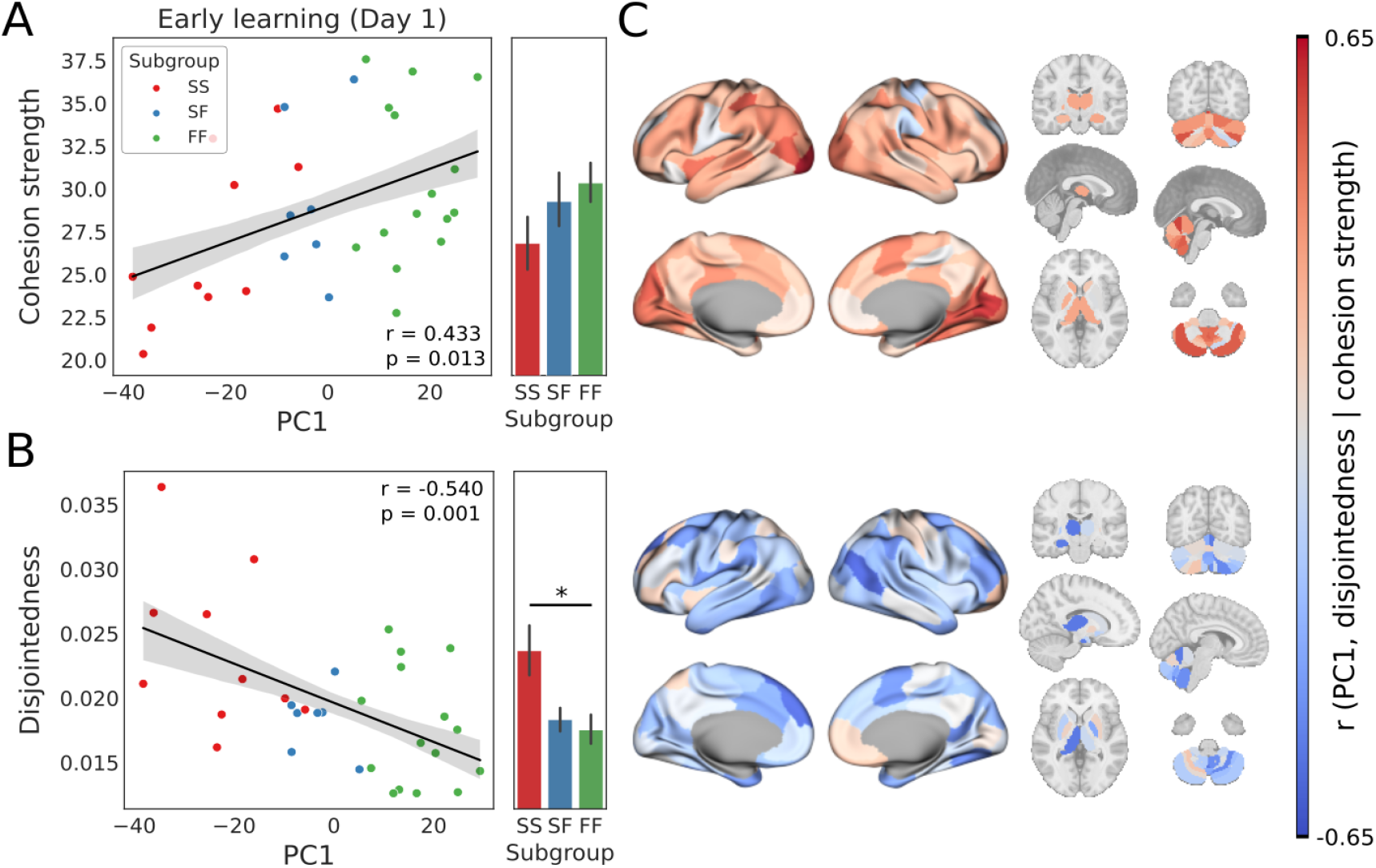
During early learning, coordinated (A) and uncoordinated (B) modular reconfiguration are associated with fast and slow learning profiles respectively. **(A)** Scatter plots show the mean strength (across regions) of cohesive flexibility (cohesion strength, see text) during early learning, plotted over PC1. Green, red and blue filled circles correspond to FF, SS and SF participants respectively. The fitted line shows the best linear fit, where the shaded area shows ±1 SE. Subgroup means ±1 SE are shown as bar graphs to the right of the scatter plots. Differences between the subgroups were non-significant [FF - SF: t(20) = 0.511, p = 0.615. FF - SS: t(23) = 1.813, p = 0.083. SF - SS: t(15) = 1.02, p = 0.324]. **(B)** Mean disjointed flexibility (disjointedness, see text) as a function of PC1 during early learning. SS was more disjointed than FF (FF > SS: t(20) = −2.855, p = 0.009), but SF did not differ significantly from SS [t(15) = −2.122, p = 0.051] or FF [t(20) = −0.435, p = 0.669]. Star in the bar plot indicates statistical significance (p < 0.05). **(C)** Brain plots show correlations between PC1 and cohesion strength (upper plots) and disjointedness (lower plots) for each region. Correlation values associated with cohesion strength (mostly positive) and disjointedness (mostly negative) are shown under a divergent colour scheme, ranging from strongly negative (dark blue) to strongly positive (dark red).

Having linked coordinated modular reconfiguration to a faster learning profile, we wondered whether *un*coordinated modular reconfiguration would be linked to a slower learning profile. We found a significant negative correlation between disjointedness during early learning and PC1 that was significantly stronger than during all other task epochs, except Day 1 baseline [1-tailed Williams test, Day 1 baseline: t(29) = −1.169, p = 0.126, adjusted p = 0.126; late learning: t(29) = −2.069, p = 0.024, adjusted p = 0.044; Day 2 baseline: t(29) = −2.171, p = 0.019, adjusted p = 0.044; early re-learning: t(29) = −1.94, p = 0.031, adjusted p = 0.044; late re-learning: t(29) = −1.877, p = 0.035, adjusted p = 0.044]. Thus, our findings not only provide evidence that uncoordinated modular dynamics are inconducive to the cognitive aspects of sensorimotor adaptation, but they also provide evidence that their effect in this regard is non-general across task epochs. As with cohesion strength above, this property appears to be a global one, as 116 out of 142 regions showed a negative correlation between PC1 and disjointedness, 27 of which (19%) were significant (−0.595 < r < −0.356, 3.255e-4 < p < 0.046). In contrast, none of the remaining 26 positive correlations with PC1 were significant (0.003 > r > 0.208, 0.253 < p < 0.984).

### Greater recruitment of a network of higher-order brain regions during early learning is associated with faster learning

Having determined that participants’ whole-brain functional networks showed significant modularity, and that cohesion strength during early learning was associated with a fast learning profile, we asked whether the recruitment of any specific brain module would be linked to fast learning. To this end, we first sought to characterise the dynamic modules (brain subnetworks) undergoing reconfiguration during early learning, calculating the proportion of all modular partitions (over all participants and time windows) in which each pair of regions was placed in the same module, referred to as a module allegiance matrix (Bassett et al. 2015). To identify specific brain networks present across time, we clustered this matrix with symmetric non-negative matrix factorisation (SymNMF, see Methods). This clustering effectively identifies a consensus partition, or set of networks that summarises the composition of the dynamic modules (Bassett et al. 2013). We took this data-driven approach to network identification because there is a systematic relationship between the networks derived from the module allegiance matrix and the temporal modules from which this matrix is calculated, which would not be the case with pre-identified resting-state networks. The summary networks and the module allegiance matrix from which they were derived are shown in Figure 4.

**Figure 4.**
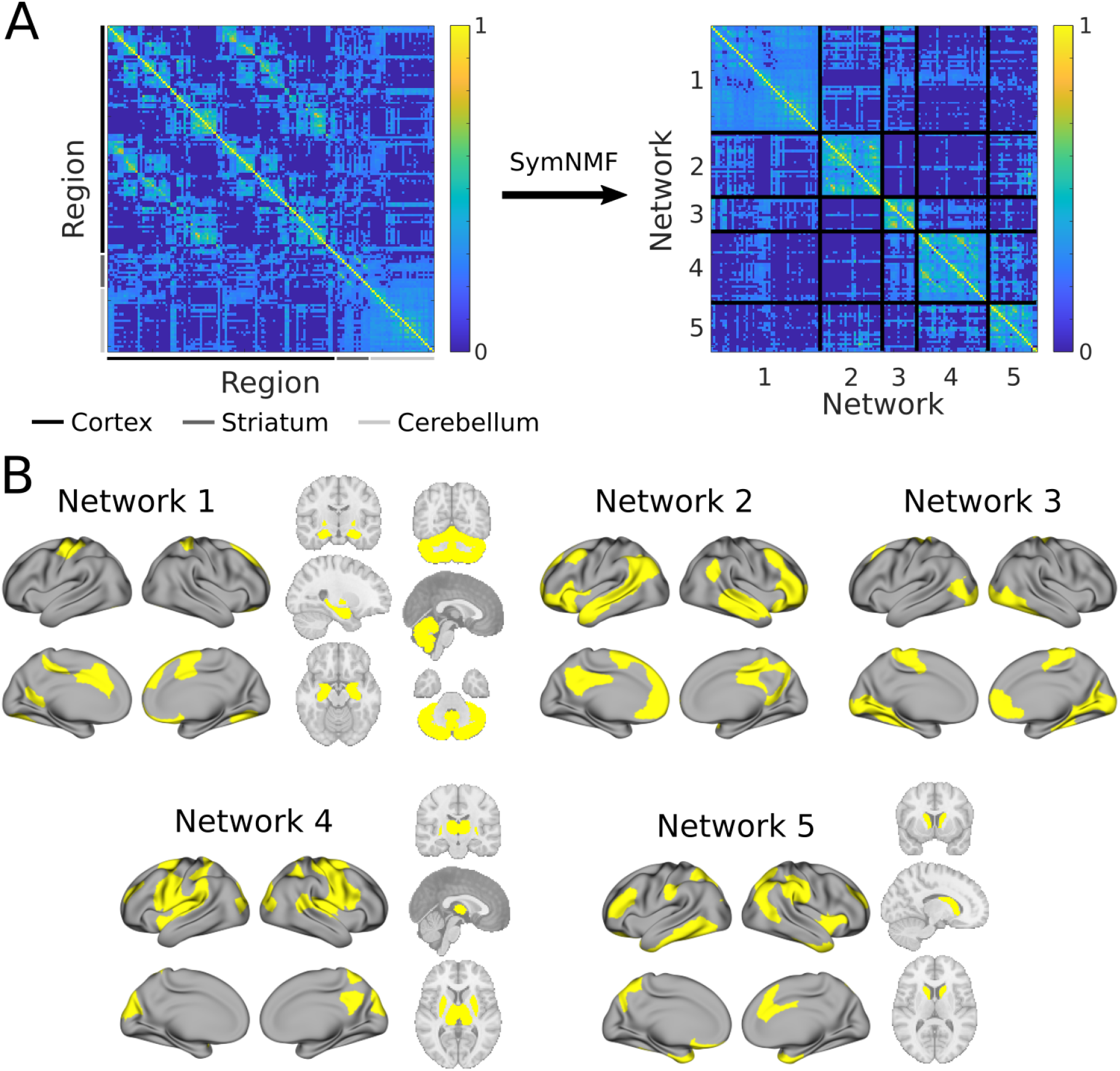
Summary of dynamic network architecture during sensorimotor adaptation. **(A)** Left, module allegiance matrix showing the probability that each pair of brain regions was in the same temporal module during early learning, calculated over all participants, modular partitions and time windows. Right, the same matrix organised according to the consensus clustering solution (see text). **(B)** Network clusters from ‘A’, rendered onto the cortical and subcortical surfaces (areas in yellow denote the derived networks). Network 1 consisted of regions spanning contralateral motor cortex, bilateral cerebellum, medial prefrontal cortex and several subcortical structures (bilateral hippocampus, pallidum, amygdala and accumbens). Network 2 was a purely cortical network, consisting of regions spanning angular gyrus, superior temporal gyrus, cingulate cortex and medial prefrontal cortex. Network 3 was also a purely cortical network, consisting mainly of regions in visual and fusiform cortex, and medial somatomotor cortex. Network 4 consisted of regions in visual cortex, medial and lateral parietal cortex, lateral somatomotor and premotor cortex, along with bilateral thalamus and putamen. Lastly, Network 5 consisted of regions spanning the anterior temporal pole, inferior and superior parietal, dorsolateral prefrontal cortex and the bilateral caudate. Note that we refrain from linking these specific task-derived networks to those described in the resting-state literature (Yeo et al. 2011) and refer to them as Networks 1 to 5 in the text. For the list of specific brain regions belonging to each network, see Supplementary Table S1.

Next, we sought to quantify the degree of regional interaction within and between the different summary networks over time. With each brain region (for each participant) assigned to one of the five networks, the interaction between any two networks can be measured by *I*_*k*1,*k*2_ = (∑*i* ∈ *C*_*k*1_, *j* ∈ *C*_*k*2_ *P*_*i,j*_.)/(|*C*_*k*1_, ||C_*k*2_|) (Bassett et al. 2015), where *C*_*k*∈1,2_are modules, |*C*_*k*_| is the number of regions they contain and *P*_*i,j*_ is the module allegiance matrix (described above; Figure 4A) capturing the proportion of time windows in which regions *i* and *j* were in the same module. We refer to the interaction of a network with itself as recruitment (Bassett et al. 2015), calculated by allowing *k*1 = *k*2. The integration between two networks *k*1 ≠ *k*2 is the normalised interaction between them 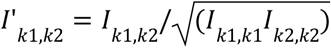, referred to as network integration (Bassett et al. 2015). In effect, recruitment captures the stability of a network’s membership over time, whereas integration captures the interactions between networks over time. An analysis of variance showed a strong effect of network on each measure (recruitment: F(4)=25.07, p=5.039e-16; integration: F(9)=13.69 p=1.642e-18), indicating that the magnitude of recruitment and integration differed across networks and their interactions.

To determine if any of the networks were selectively associated with participants’ learning profiles, we calculated the Pearson correlation coefficient between PC1 and recruitment of each network during early learning, correcting for multiple comparisons across the five networks (see Methods). We found a significant positive correlation for Network 5 (Pearson’s r = 0.557, p = 9.227e-4, adjusted p = 0.005), a network composed mainly of anterior temporal pole and prefrontal regions, in which recruitment was statistically indistinguishable between the FF and SF subgroups, and was significantly higher in FF and SF than SS (Figure 5A). The composition of Network 5 (including regions typically associated with higher-order cognitive processes, *e*.*g*. limbic cortex) is consistent with the hypothesis that FF and SF participants were using cognitive strategies during learning on Day 1.

**Figure 5.**
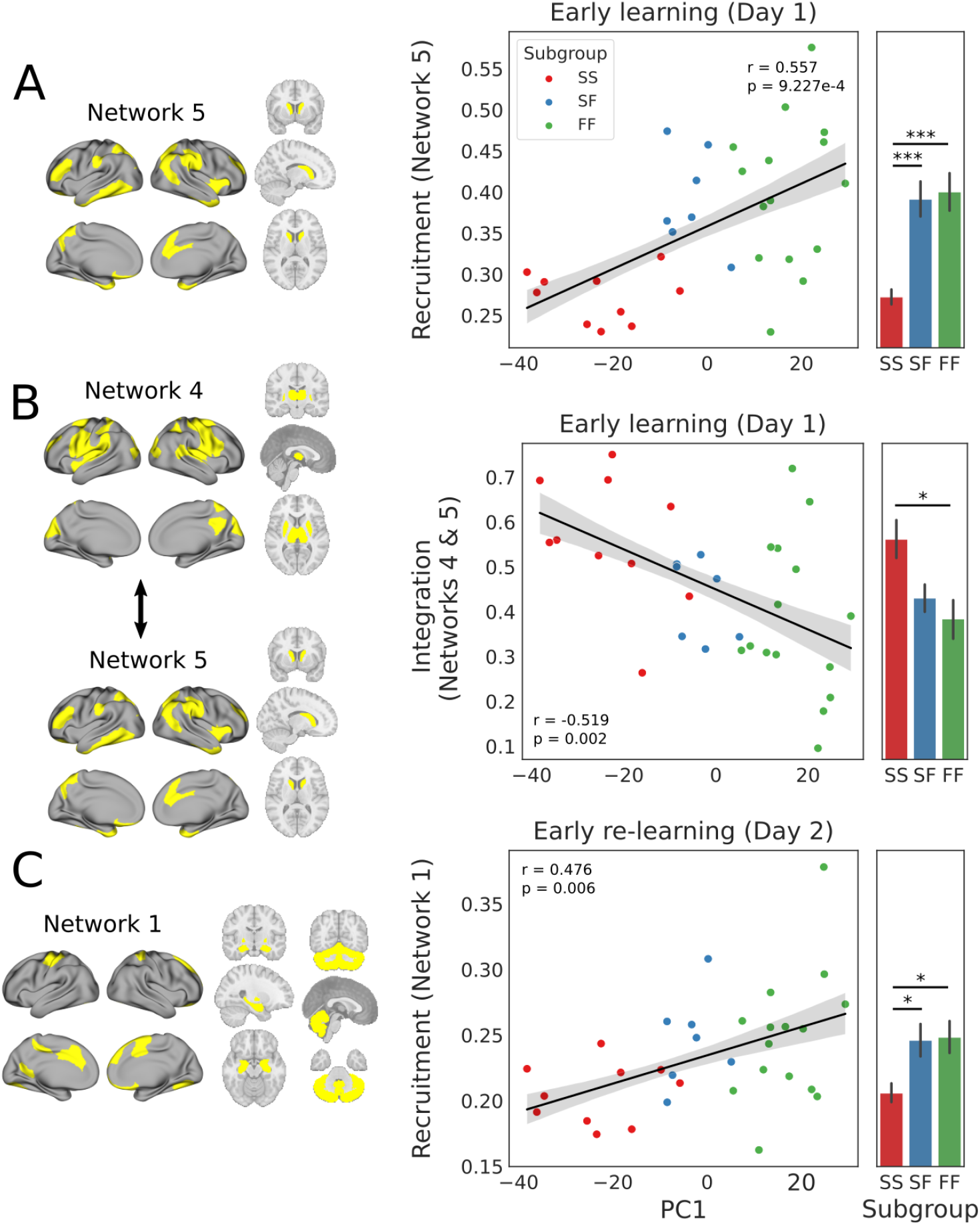
Recruitment of specific summary networks is associated with participants’ learning profiles during early learning (A) and re-learning (C) on each day. **(A)** During early learning, recruitment of Network 5 (composed mainly of anterior temporal and prefrontal regions, left side) was positively correlated with PC1, where recruitment by FF and SF participants was statistically indistinguishable [2-sample t-test, t(20) = 0.235, p = 0.817] but was greater among FF [t(23) = 4.273, p = 2.85e-4] and SF [t(15) = 5.398, p = 7.398e-5] than SS (rightmost panel). **(B)** During early learning, integration between Network 4 (composed mainly of visual and somatomotor regions, left side) and Network 5 was negatively correlated with PC1, where integration between these networks was greater among FF participants than SS [t(23) = −2.654, p = 0.014] but did not differ statistically between FF and SF [t(20) = −0.652, p = 0.522] nor between SF and SS [t(20) = −2.13, p = 5.016e-2] (rightmost panel). **(C)** During early re-learning, recruitment of Network 1 (composed mainly of hippocampal, striatal and cerebellar regions, left side) was positively correlated with PC1, where recruitment by FF and SF participants was statistically indistinguishable [t(20) = 0.11, p = 0.913] but was greater among FF [t(23) = 2.492, p = 0.02] and SF [t(15) = 2.86, p = 0.012] participants than SS (rightmost panel). In scatter plots, fitted line shows linear fit, where shading corresponds to ±1 SE. Bar plots show means, where error bars show ±1 SE. In bar plots, stars indicate significant differences (one star: p < 0.05; three stars: p < 1e-3).

To determine whether the correlation between PC1 and the recruitment of Network 5 was strongest during early learning, we compared it to the same correlation during the other epochs, finding no significant differences [1-tailed Williams test, Day 1 baseline: t(29) = 1.462, p = 0.077, adjusted p = 0.232; late learning: t(29) = 1.037, p = 0.139, adjusted p = 0.232; Day 2 baseline: t(29) = −0.255, p = 0.600, adjusted p = 0.600; early re-learning: t(29) = 1.210, p = 0.118, adjusted p = 0.232; late re-learning: t(29) = 0.183, p = 0.428, adjusted p = 0.535]. Thus, this brain-behaviour relationship was not learning-specific.

Next, we sought to determine whether the integration of specific pairs of networks had bearing on participants’ learning profiles. We therefore calculated the integration of all (10 = 5 × 4 / 2) pairs of networks during early learning, and calculated the Pearson correlation between each integration measure and PC1, correcting for multiple comparisons. There was a significant negative correlation between PC1 and the integration of Network 5 with a visuo-somatomotor network (Network 4, Figure 5B). To determine if this correlation was most pronounced during early learning, we tested whether it was stronger than during the other epochs, finding that it was significantly stronger than during all other epochs except late learning and Day 2 baseline [1-tailed Williams test, Day 1 baseline: t(29) = −3.761, p = 3.810e-4, q = 0.002; late learning: t(29) = −1.092, p = 0.142, adjusted p = 0.142; Day 2 baseline: t(29) = −1.400, p = 0.086, adjusted p = 0.108; early re-learning: t(29) = −2.467, p = 0.010, adjusted p = 0.019; late re-learning: t(29) = −2.405, p = 0.011, adjusted p = 0.019]. Thus, while the recruitment of Network 5 during early learning was predictive of a fast learning profile that emerged across days, the propensity of this network to support fast learning appears to have depended on its relative segregation from Network 4. Given the regional composition of Networks 4 and 5, this finding suggests that the successful implementation of a cognitive strategy not only requires the recruitment of a suitable higher-order brain network (putatively Network 5), but also requires that this network be decoupled from a sensorimotor network that (presumably) implements the visual-to-motor mapping.

### Greater recruitment of a hippocampal-striatal-cerebellar network during early re-learning is associated with a faster learning profile

Having associated a fast learning profile with the recruitment of a network of higher order brain regions during early learning (Network 5), we sought to determine whether this same network would be associated with re-learning on the second day, or alternatively, whether another network (or networks) would be associated with participants’ learning profiles. To this end, we calculated the Pearson correlation coefficient between PC1 and the recruitment of each network during early re-learning, correcting for multiple comparisons across the five networks. Note that we used the above networks for this analysis (derived during early learning and shown in Figure 4) but our results were qualitatively unchanged when we derived the summary networks during early re-learning. Indeed, our Day 1 results were also qualitatively unchanged (see the Supplementary Materials). We did not find a significant correlation for Network 5, but we did find a significant positive correlation for a network composed mainly of subcortical regions, including bilateral hippocampus, amygdala, accumbens, pallidum and the entire cerebellum (Network 1, Figure 5C). Comparison of this correlation to that during the other five epochs showed that this brain-behaviour relationship was stronger on Day 2 [1-tailed Williams test, Day 1 baseline: t(29) = 1.958, p = 0.030, adjusted p = 4.998e-2; early learning: t(29) = 2.924, p = 0.003, adjusted p = 0.008; late learning: t(29) = 3.770, p = 3.724e-4, adjusted p = 0.002; Day 2 baseline: t(29) = 0.833, p = 0.206, adjusted p = 0.257; late re-learning: t(29) = 0.353, p = 0.363, adjusted p = 0.363]. Furthermore, the recruitment of Network 1 during early re-learning by the FF and SF subgroups was statistically indistinguishable, but was significantly higher in FF and SF than SS (Figure 5C, right panel). Under the assumption that the hippocampus is involved in the recall of an explicit cognitive strategy during adaptation (Haith, Huberdeau, and Krakauer 2015; McDougle et al. 2022; de Brouwer et al. 2021, 2018) this finding provides evidence that the FF and SF subgroups not only approached the task in the same (or a similar) way on Day 2, but also on Day 1. The composition of Network 1 is also consistent with emerging evidence for direct neuroanatomical projections (Bostan and Strick 2018; Bostan, Dum, and Strick 2018) and functional interactions (Wagner et al. 2017; Larry et al. 2019; Kostadinov et al. 2019) between reward-related striatal circuits and the cerebellum, as well as the more general role of the cerebellum in higher-order cognitive processes (Marek et al. 2018).

## Discussion

Humans show marked variability in their capacity for sensorimotor adaptation, the neural bases of which are unclear. We characterised this variation and investigated its whole-brain correlates by having participants perform a sensorimotor adaptation task on two testing days, separated by 24 hrs. A clustering of participants’ learning data across both days revealed three distinct profiles of learners: participants who exhibited fast learning on both days (the FF subgroup), suggesting the use of explicit, cognitive strategies toward the task (de Brouwer et al. 2018; Brouwer et al., n.d.); participants who exhibited slow learning on both days (the SS subgroup), suggesting primarily implicit learning (de Brouwer et al. 2018); and participants who exhibited slow learning on Day 1, but fast learning on Day 2 (the SF subgroup) (Figure 2D,E). Participants’ whole-brain functional networks showed significant modularity during the task, where more coordinated modular reconfiguration during early learning was linked to faster learning (Figure 3), as was the recruitment of a functional network involving regions in anterior temporal and prefrontal cortex (Network 5, Figure 5A). Notably, greater recruitment of this network distinguished the FF and SF subgroups from the SS subgroup, and greater integration of Network 5 with a visuo-somatomotor network (Network 4) was associated with slower learning (Figure 5B). Thus, the propensity to segregate functional networks associated with (presumably) the cognitive and sensorimotor aspects of the task offers a potential explanation for the differences in behavioural performance by the subgroups. Finally, we found that greater recruitment of a large network involving hippocampus, ventral striatum and cerebellum (Network 1) during early re-learning was also linked to faster learning (Figure 5C and Figure 6), and distinguished the FF and SF subgroups from SS (Figure 5C). Under the hypothesis that savings reflects the recall of a previously learned strategy (Haith, Huberdeau, and Krakauer 2015; Morehead et al. 2015) this neural finding not only provides evidence that FF and SF took a similar approach to re-learning, but by extension, it implies that SF participants had found an appropriate re-aiming strategy by the end of Day 1. Taken together, our findings provide a novel characterisation of both behaviour and whole-brain dynamics during human sensorimotor adaptation, and suggest that the selective recruitment and isolation of cognitive systems during early learning supports the implementation of explicit strategies during adaptation.

**Figure 6.**
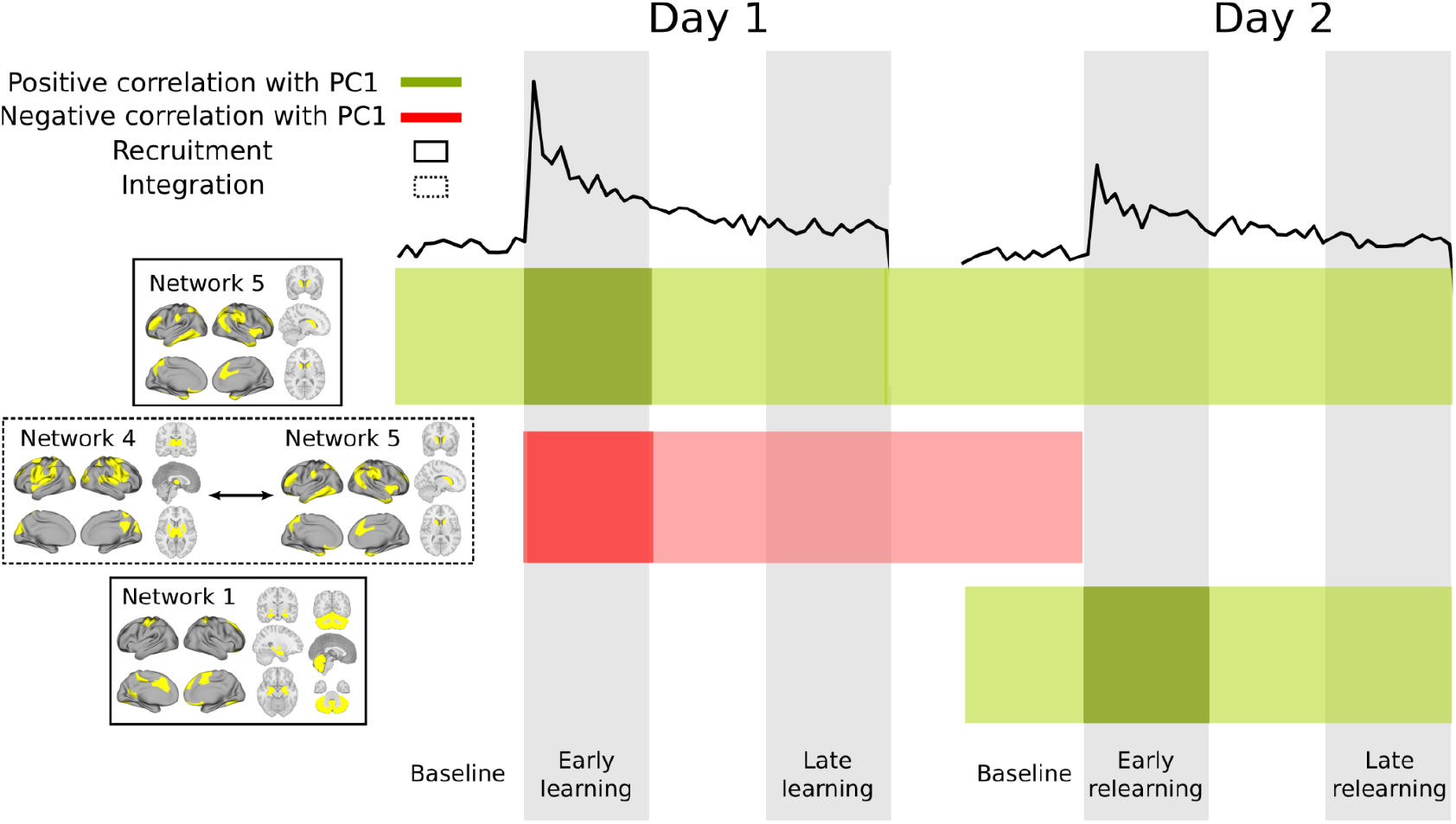
Summary of network recruitment and integration during learning and re-learning. Recruitment of Network 5 (left, top solid box) during early learning was correlated with a fast learning profile (dark green shading) but this correlation did not differ statistically from that during other task epochs on either day (light green). Integration of Networks 4 and 5 (left, dotted box) was correlated with a slow learning profile (dark red shading). This effect was stronger than during Day 1 baseline, and early and late re-learning (light red). Recruitment of Network 1 (left, bottom solid box) during early re-learning was correlated with a fast learning profile. This effect was specific to Day 2. Black traces at top show group-averaged error.

### A convergence of regional-level and network-level approaches

Prior neuroimaging studies of sensorimotor adaptation in humans have documented changes in activation during learning in a broad range of brain regions, including prefrontal, cingulate, premotor, motor and parietal cortices, as well as in the striatum and cerebellum (R. Shadmehr 1997; Anguera et al. 2007; Doyon et al. 2009; Dayan and Cohen 2011; Krakauer et al. 2004; S. Kim et al. 2015; Hardwick et al. 2013; Bédard and Sanes 2014). Identifying the specific roles of these regions has been challenging (Dayan and Cohen 2011; Hardwick et al. 2013) but a key theme to emerge from this work is that the early and late phases of adaptation appear to engage different neural systems. For example, studies have shown increases in the activation of several higher-order brain regions during early learning, such as dorsolateral prefrontal cortex (DLPFC) and anterior cingulate cortex (Anguera et al. 2010, 2011), where differential activation appears in other regions during late learning, such as the cerebellum (Seidler and Noll 2008; S. Kim et al. 2015; Flament et al. 1996). Additionally, better visuospatial working memory (Christou et al. 2016) and stronger activation of DLPFC (Anguera et al. 2010) have been shown to correlate with faster adaptation across participants. These findings suggest an important role for cognitive brain systems during the early stages of learning in particular, and may explain why cognitive deficits associated with ageing and explicit memory have been linked to slower adaptation (Anguera et al. 2011; Trewartha et al. 2014; Wolpe et al. 2020).

Our results are generally consistent with these observations, but they extend this body of work to whole-brain networks and their dynamics, and provide evidence that faster learners are distinguished by their ability to put together (recruit) functional networks to suit the cognitive aspect of sensorimotor adaptation. We propose that this ability corresponds to strategic thinking more generally (Brouwer et al., n.d.) and that the coordinated reconfiguration of functional networks is its neural signature. Future work is required to investigate this proposition, but it is noteworthy that cohesion strength and the recruitment of Network 5 were both correlated with PC1 during early learning, when cognitive strategies are believed to be most influential on task performance (de Brouwer et al. 2018; Taylor, Krakauer, and Ivry 2014), and that cohesion strength correlates negatively with the depth of unconsciousness (Standage et al. 2020), a state that is reasonably described as cognitively disengaged.

### Methodological considerations and limitations

The above descriptions of individual differences in modular dynamics offer a novel characterisation of the (possible) whole-brain bases of the explicit component of sensorimotor adaptation, but they do not speak to *how* these dynamics may support the explicit component. The time-specificity of our results may be instructive in this regard (Figure 6). The correlation between cohesion strength and PC1 was not specific to early learning, as this correlation did not significantly differ from that during other epochs of the two day task, nor did the recruitment of Network 5 (Figure 6). Thus, these brain-behaviour relationships reveal properties of brain dynamics and network composition that appear to be conducive to the formation and/or use of cognitive strategies, but that do not directly correspond to aspects of the learning process, *e*.*g*. the search for a strategy. This finding is consistent with earlier behavioural analyses providing evidence that sensorimotor adaptation is never (on the timescale of most experiments) entirely explicit or implicit (Taylor, Krakauer, and Ivry 2014; de Brouwer et al. 2018; Huberdeau, Krakauer, and Haith 2019) and with the more general notion that humans do not turn ‘cognition’ on and off like a switch. In this regard, our finding that the integration of a sensorimotor network (Network 4) with a network involving higher-order association areas (Network 5) was detrimental to fast learning may also be instructive, since this effect was not the case during Day 1 baseline or re-learning. As such, it appears that fast learning involves the ability to segregate the functional networks recruited for task performance, in conjunction with the above more general properties of cognition. Our study does not speak to how participants implement this segregation, nor its reliance on whole-brain dynamics. Large-scale neural modelling techniques offer a principled approach for asking such questions (Shen et al. 2019; Honey et al. 2009) and future work should leverage these methods in the study of sensorimotor adaptation.

Our behavioural analysis leveraged the variation in participants’ learning rates to infer the relative contribution of the explicit component to task performance, rather than directly estimating its contribution by having participants report their aiming directions during the task (Taylor, Krakauer, and Ivry 2014). The latter approach may have allowed us to make stronger claims about the links between cognitive strategies and brain networks, but ‘reporting trials’ have been shown to magnify (and thereby distort) the use of such strategies and their inclusion would presumably have changed the nature of the continuous BOLD data collected, impacting our construction and interpretation of temporal networks. Nevertheless, given the well-established correlation between participants’ aiming directions and their rates of learning (Brouwer et al., n.d.; de Brouwer et al. 2018) it is reasonable to use participants’ level of adaptation during early learning to index the magnitude of the explicit component.

We have presented converging behavioural (Figure 2D) and neural (Figure 5A, C) evidence that the SF subgroup was unsuccessfully attempting a cognitive strategy during early learning on Day 1, but this evidence does not speak to why SF may have been unsuccessful at that time. To a degree, our use of PC1 as a proxy for group membership limits our ability to identify neural measures that differentiate these two subgroups, since the magnitude of neural measures that correlate significantly with PC1 will tend to occur along a gradient from SS to FF (or vice versa) with SF in between. Suffice to say, the neural measures used in our study were sufficient to demonstrate similarities in whole-brain dynamics by the FF and SF subgroups, but were insufficient to demonstrate whole-brain differences that may have explained the stark differences in their Day 1 behaviour. There are many possible approaches to addressing this limitation, such as deriving subgroup-specific summary networks and investigating their differences, but these approaches are beyond the scope of the present work.

### Neural evidence for savings as the recall of a cognitive strategy

One of the striking features of our behavioural clusters is the extent to which savings differed across subgroups. Savings in visuomotor rotation tasks are widely believed to be driven by the recall of a successful re-aiming strategy, which has been directly linked to the explicit component (Haith, Huberdeau, and Krakauer 2015; Morehead et al. 2015). If so, then faster learners might be expected to express greater savings (de Brouwer et al. 2018). Here, we found that fast learners on Day 1 did indeed show significant savings (the FF subgroup). However, we found that the largest savings (∼3 times larger than FF) were expressed by the SF subgroup (Figure 2D), who were actually slow learners on Day 1. At first glance, this finding may not conform to expectations, but it is nevertheless intuitive in light of how savings is calculated. Fast learners, by definition, have limited scope for savings, since their rates of adaptation are already fast and are therefore difficult to improve. In contrast, slow learners have ample scope for improvement when they re-encounter the rotation on Day 2. As such, ideal candidates for savings are participants who identify a re-aiming strategy late on Day 1, giving them both scope for improvement and a successful strategy to be recalled.

A striking neural finding from our study was that the FF and SF subgroups could not be distinguished by their recruitment of Network 5 during early learning or their recruitment of Network 1 during early re-learning, whereas both subgroups recruited each of these networks more strongly than the SS subgroup. The composition of Network 5 — including regions in limbic and prefrontal cortex — is clearly suggestive of a role in explicit processes, such as the explicit contribution to visuomotor adaptation (Figure 4B). In contrast, the composition of Network 1 warrants further comment in relation to recall and savings. Given the extensive body of evidence for hippocampal involvement in declarative memory (Eichenbaum 2004; Duff et al. 2019; Squire 2004), the inclusion of bilateral hippocampus in Network 1 is clearly consistent with the hypothesis that it supports the recall of a successful re-aiming strategy (Haith, Huberdeau, and Krakauer 2015; McDougle et al. 2022; de Brouwer et al. 2018; Brouwer et al., n.d.; Morehead et al. 2015). In turn, this hypothesis is consistent with our finding that the FF and SF subgroups both showed significant savings (Figure 2D) and both recruited Network 1 more strongly than the SS subgroup (Figure 5C). However, the inclusion of bilateral cerebellum in Network 1 may be considered unconventional, given the extensive body of evidence for its involvement in the *implicit* component of sensorimotor adaptation (Martin et al. 1996; Tseng et al. 2007; Izawa, Criscimagna-Hemminger, and Shadmehr 2012) and in motor learning and control more generally (Wolpert, Miall, and Kawato 1998; Reza Shadmehr, Smith, and Krakauer 2010; Reza Shadmehr and Krakauer 2008). This traditional characterisation of cerebellar function, however, is evolving in light of recent evidence for its role in more domain-general and cognitive functions (King et al. 2019; Diedrichsen et al. 2019; Sokolov, Miall, and Ivry 2017). Indeed, recent functional mapping studies suggest that a much larger extent of the cerebellum is associated with the default mode and fronto-parietal resting-state networks, which are extensively correlated with higher-order cognitive processes, than with somatomotor resting-state networks (Marek et al. 2018). This body of work supports the current consensus that the cerebellum plays an important role in human cognition more generally, and that its processes are not just motor-related (Strick, Dum, and Fiez 2009; Buckner 2013; Buckner et al. 2011; Stoodley and Schmahmann 2009; Diedrichsen et al. 2019; King et al. 2019). We can only speculate on the nature of cerebellar involvement during re-learning on our task but our findings suggest a role for distributed subcortical systems (hippocampus, striatum and cerebellum) in the consolidation and recall of strategic learning processes, or alternatively, competitive and/or compensatory interactions between parallel learning systems (Albert et al. 2022; Miyamoto, Wang, and Smith 2020). The latter possibility is particularly intriguing in light of the recent finding that explicit re-learning (supporting savings) coincides with attenuated implicit adaptation (Avraham et al. 2021). As such, our Network 1 may reflect functional interactions between explicit systems supporting recall (e.g. hippocampus) and implicit learning (e.g. cerebellum), governing the changing balance of their respective contributions to task performance.

## Conclusion

We have characterised three behavioural profiles of sensorimotor adaptation that emerge across two days of learning and re-learning, and we have linked these profiles to the dynamics and composition of whole-brain networks during early learning, when cognitive strategies are believed to be most prominent on this class of task (de Brouwer et al. 2018; Taylor, Krakauer, and Ivry 2014). Our findings suggest that coordinated modular reconfiguration supports the recruitment of functional networks to suit the task at hand, where fast learning requires a decoupling of networks associated with cognitive and sensorimotor aspects of task performance, and fast re-learning involves a functional coupling of hippocampal and cerebellar systems. The latter may support compensatory interactions between explicit and implicit learning systems and their respective contributions to performance. Overall, our findings support the hypothesis that whole-brain dynamics of cognition drive individual differences in learning.

## Methods

### Experimental Design and Statistical Analysis

40 right-handed individuals between the ages of 18 and 35 (M = 22.5, SD = 4.51; 13 males) participated in the study and received financial compensation for their time. Data from 8 participants were excluded, due to either head motion in the MRI scanner (N=4; motion greater than 2 mm or 2° rotation within a single scan) or their inability to learn the rotation task (N=4), leaving 32 participants in the final analysis. Right-handedness was assessed with the Edinburgh handedness questionnaire (Oldfield 1971). Participants’ written, informed consent was obtained before commencement of the experimental protocol. The Queen’s University Research Ethics Board approved the study and it was conducted in accordance with the principles outlined in the Canadian Tri-Council Policy Statement on Ethical Conduct for Research Involving Humans and the principles of the Declaration of Helsinki (1964).

Experimenters were not blind to testing. Statistical significance was defined by alpha pre-set to 0.05. Error bars indicate +/- 1 standard errors of the mean (SEM). Statistical tests are described in the figure legends and each test was selected based on data distributions using histograms. Two-tailed tests were used unless otherwise noted. Differences between correlations were tested for significance using Williams’ method (Steiger 1980). False discovery rate (FDR) corrections for family-wise error rate used the method by Benjanimi and Hochberg (1995).

### Apparatus

In the scanner, participants performed hand movements that were directed towards a target by applying a directional force onto an MRI-compatible force sensor (ATI technologies) using their right index finger and thumb. The target and cursor stimuli were rear-projected with an LCD projector (NEC LT265 DLP projector, 1024 × 768 resolution, 60 Hz refresh rate) onto a screen mounted behind the participant. The stimuli on the screen were viewed through a mirror fixed to the MRI coil directly above participants’ eyes, thus preventing participants from being able to see the hand. The force sensor and associated cursor movement were sampled at 500 Hz.

### Procedure

This experiment used a well-established visuomotor rotation task (Krakauer 2009) to probe sensorimotor adaptation (Figure 1*A*). To start each trial, the cursor (20-pixel radius) appeared in a central start position (25-pixel radius). A white target circle (30-pixel radius) presented on a black screen appeared at one of eight locations (0, 45, 90, 135, 180, 225, 270, 315º) on an invisible ring around the central position (300-pixel radius) and filled in (white) following a 200 ms delay. Once filled in, participants applied a brief directional force impulse on the force sensor (threshold of 1.5 N), which launched the cursor towards the target. Targets could randomly appear in each location, without replacement, during sets of 8 trials. Once launched, the cursor would travel the 300-pixel distance to the ring over a 750 ms period (with a bell-shaped velocity profile) before becoming stationary at the ring to provide participants with end-point error feedback. If the cursor overlapped with the target to any extent, the target would become green, signifying a target ‘hit’. Each trial was separated by 4 s and within this period, participants had 2.6 s from target presentation to complete the trial (including the 200 ms target delay, participants’ own reaction time, and the 750 ms cursor movement; any remaining time was allotted to providing the end-point error feedback). At 2.6 s the trial was ended, the screen was blanked, the data saved, and participants would briefly wait for the next trial to begin. Reaction times were not stressed in this experimental procedure. On trials in which the reaction time exceeded 2.6 s, the trial would end, and the participant would wait for the next trial to begin. These discarded trials were rare (0.56% across all trials, all participants) and were excluded from behavioural analyses, but were kept in the neuroimaging analysis due to the continuous nature of the fMRI task and our focus on functional connectivity analyses.

During each testing session, 120 ‘baseline’ trials (15 sets of 8 trials) were completed without a rotation of the cursor. Following these trials, 320 ‘learning’ trials (40 sets of 8 trials) were completed, wherein a 45º clockwise rotation of the cursor was applied. The baseline and learning trials were completed during one continuous fMRI scan. Following this scan, conditions were restored to baseline (i.e. no rotation of cursor) in a separate scan and participants performed 120 ‘washout’ trials. These washout trials allowed us to probe participants’ rate of relearning 24 hours later (and thus, their savings). In addition to these VMR-related task components, we also interspersed three 6-minute resting state fMRI scans prior to, between, and following VMR learning and washout (note that spontaneous brain activity during these resting scans is not considered in this paper). During these resting-state scans, participants were instructed to rest with their eyes open, while fixating a central cross location presented on the screen. The total testing time was 75 minutes on each testing day.

### MRI Acquisition

Participants were scanned using a 3-Tesla Siemens TIM MAGNETOM Trio MRI scanner located at the Centre for Neuroscience Studies, Queen’s University (Kingston, Ontario, Canada). Functional MRI volumes were acquired using a 32-channel head coil and a T2*-weighted single-shot gradient-echo echo-planar imaging (EPI) acquisition sequence (time to repetition (TR) = 2000 ms, slice thickness = 4 mm, in-plane resolution = 3 mm × 3 mm, time to echo (TE) = 30 ms, field of view = 240 mm × 240 mm, matrix size = 80 × 80, flipb angle = 90°, and acceleration factor (integrated parallel acquisition technologies, iPAT) = 2 with generalised auto-calibrating partially parallel acquisitions (GRAPPA) reconstruction. Each volume comprised 34 contiguous (no gap) oblique slices acquired at a ∼30° caudal tilt with respect to the plane of the anterior and posterior commissure (AC-PC), providing whole-brain coverage of the cerebrum and cerebellum. A T1-weighted ADNI MPRAGE anatomical was also collected (TR = 1760 ms, TE = 2.98 ms, field of view = 192 mm × 240 mm × 256 mm, matrix size = 192 × 240 × 256, flip angle = 9°, 1 mm isotropic voxels). For each resting-state scan, 180 imaging volumes were collected. For the baseline and learning epochs, a single, continuous scan was collected of 896 imaging volumes duration. For the washout scan, one scan of 256 imaging volumes was collected. Each scan included an additional 8 imaging volumes at both the beginning and end of the scan.

On Day 1, a separate practice session was carried out before the actual fMRI experiment to familiarise participants with the apparatus and task. This session involved performing 60 practice baseline (non-rotation) trials. The fMRI testing session for each participant lasted 2 hours and included set-up time (∼20 min.), practice (∼10 min.), one high-resolution anatomical scan (∼8 min duration), two DTI scans (one in the AP direction and the other in the PA direction; ∼10 min duration), a resting-state scan (6 min duration), a baseline and rotation scan (∼30 min duration), a resting-state scan (6 min. duration), a washout scan (∼9 min duration), and a final resting-state scan (6 min. duration). In the present paper, we only consider results associated with the baseline-rotation scan collected on Day 1.

### MRI Preprocessing

Preprocessing was performed using fMRIPrep 1.4.0 (Esteban, Ciric, et al. 2019; Esteban et al. 2018) RRID:SCR_016216), which is based on Nipype 1.2.0 (Esteban, Markiewicz, et al. 2019); RRID:SCR_002502).

#### Anatomical data preprocessing

T1w scans in each session were corrected for intensity non-uniformity (INU) with N4BiasFieldCorrection (Tustison et al. 2010), distributed with ANTs 2.2.0 (Avants et al. 2008) RRID:SCR_004757). The T1w-reference was then skull-stripped with a Nipype implementation of the antsBrainExtraction.sh workflow (from ANTs), using OASIS30ANTs as target template. Brain tissue segmentation of cerebrospinal fluid (CSF), white-matter (WM) and grey matter (GM) was performed on the brain-extracted T1w using fast (FSL 5.0.9, RRID:SCR_002823, (Zhang, Brady, and Smith 2001). A T1w-reference map was computed after registration of the individual T1w scans (after INU-correction) using mri_robust_template (FreeSurfer 6.0.1, (Reuter, Rosas, and Fischl 2010). Brain surfaces were reconstructed using recon-all (FreeSurfer 6.0.1, RRID:SCR_001847, (Dale, Fischl, and Sereno 1999), and the brain mask estimated previously was refined with a custom variation of the method to reconcile ANTs-derived and FreeSurfer-derived segmentations of the cortical grey matter of Mindboggle (RRID:SCR_002438, (Klein et al. 2017). Volume-based spatial normalisation to FSL’s MNI ICBM 152 non-linear 6th Generation Asymmetric Average Brain Stereotaxic Registration Model ((Evans et al. 2012), RRID:SCR_002823; TemplateFlow ID: MNI152NLin6Asym) was performed through nonlinear registration with antsRegistration (ANTs 2.2.0), using brain-extracted versions of both T1w reference and the T1w template.

#### Functional data preprocessing

For each of the BOLD runs per participant, the following preprocessing was performed. First, a reference volume and its skull-stripped version were generated using a custom methodology of fMRIPrep. The BOLD reference was then co-registered to the T1w reference using bbregister (FreeSurfer) which implements boundary-based registration (Greve and Fischl 2009). Co-registration was configured with nine degrees of freedom to account for distortions remaining in the BOLD reference. Head-motion parameters with respect to the BOLD reference (transformation matrices, and six corresponding rotation and translation parameters) are estimated before any spatiotemporal filtering using mcflirt (FSL 5.0.9, (Jenkinson et al. 2002). BOLD runs were slice-time corrected using 3dTshift from AFNI 20160207 ((Cox and Hyde 1997), RRID:SCR_005927). The BOLD time-series were resampled into standard space (MNI152NLin6Asym), correspondingly generating spatially-normalised, preprocessed BOLD runs. First, a reference volume and its skull-stripped version were generated using a custom methodology of fMRIPrep. Several confounding time-series were calculated based on the preprocessed BOLD. The six head-motion estimates calculated in the correction step were included as confounding time series, along with the temporal derivatives and quadratic terms for each. A set of physiological regressors were extracted to allow for component-based noise correction (Behzadi et al. 2007). Here, principal components are estimated after high-pass filtering the preprocessed BOLD time-series (using a discrete cosine filter with 128s cut-off) based on anatomical masks (aCompCor). We used aCOMPCOR because it has proven to be one of the most consistent pipelines for mitigating the problem of motion across performance indices related to dynamic functional connectivity analyses (see Lydon-Staley et al., 2019). Components are calculated within the intersection of the aforementioned mask and the union of CSF and WM masks calculated in T1w space, after their projection to the native space of each functional run (using the inverse BOLD-to-T1w transformation). Components are also calculated separately within the WM and CSF masks. *k* components with the largest singular values are retained, and the remaining components are dropped from consideration. All resamplings were performed with a single interpolation step by composing all the pertinent transformations (i.e. head-motion transform matrices, susceptibility distortion correction when available, and co-registrations to anatomical and output spaces). Gridded (volumetric) resamplings were performed using antsApplyTransforms (ANTs), configured with Lanczos interpolation to minimise the smoothing effects of other kernels (Lanczos 1964). Non-gridded (surface) resamplings were performed using mri_vol2surf (FreeSurfer).

Many internal operations of fMRIPrep use Nilearn 0.5.2 (Abraham et al. 2014), RRID:SCR_001362), mostly within the functional processing workflow. For more details of the pipeline, see the section corresponding to workflows in fMRIPrep’s documentation.

#### Region-of-Interest (ROI) signal extraction

ROI extraction and nuisance regression was performed using Nilearn 0.5.2 (Abraham et al., 2014). For each participant, the raw BOLD time series for each voxel inside a given ROI was isolated using the Schaeffer-100 atlas for cortical areas (Schaefer et al. 2018), the Harvard-Oxford atlas for subcortical areas (Frazier et al. 2005; Desikan et al. 2006; Goldstein et al. 2007; Makris et al. 2006), and the SUIT cerebellar atlas for cerebellum (Diedrichsen et al. 2009). The mean time course of each ROI was highpass-filtered (cutoff = .01Hz), temporally detrended, and z-scored across each run in order to ensure that all voxels had the same scale of activation after preprocessing. In order to account for motion confounds and physiological noise, the above-described nuisance regressors were included. That is, the participant’s six motion parameters and their derivatives, squares, and the squares of the derivatives (24 total), and the top six aCompCor regressors. Note that nuisance regression was performed orthogonally to temporal filters so as to avoid reintroducing high-frequency noise from the regressors (see (Lindquist et al. 2019).

### Data Analysis

#### Behavioural analyses

To assess performance on the task, the angle of the cursor relative to the target at the moment the cursor reached the target distance (or nearest data point if this value could not be used) was calculated. This was done by subtracting the target angle from an angle derived from the horizontal and vertical position at a 300-pixel distance from the start point. The median endpoint cursor error for the set of 8 trials (1 trial for each target) was extracted (binned median error) and the mean binned median error for early trials (bins 1-3; 24 trials total) and late trials (bins 38-40; 24 trials total) were calculated to yield ‘early’ and ‘late’ error measures for all participants on each day. Savings for each participant was measured by subtracting the early error on Day 2 from the early error on Day 1 (Morehead et al. 2015).

To determine whether distinct subgroups of learners were present in our sample, k-means clustering (Lloyd 1982) was performed across participants using these five learning measures (early error on Day 1, early error on Day 2, late error on Day 1, late error on Day 2, and savings). We ran k-means until convergence 1000 times with random initialization for all integers in 2 ≤ k ≤ 9 (maximum of 10,000 iterations of the algorithm for each initialization). The solution with the lowest sum of Euclidean distances between the data points and their assigned centroids was chosen for each value of k, and each of these solutions was evaluated by the Silhouette (Rousseeuw 1987) and Calinski-Harabasz (Calinski and Harabasz 1974) cluster evaluation indices. The solution corresponding to k = 3 had the highest score according to both indices (see Results). Results were identical under Euclidean and Manhattan distances measures. We tested this clustering solution for statistical significance by randomly drawing from a multivariate normal distribution with the same means and covariance as the data (1000 iterations, one draw for each participant), running k-means for each iteration in the same way as above. We recorded the highest value of the Calinski-Harabasz and Silhouette indices (over all k) for each iteration, where the probability of falsely rejecting the null hypothesis (for each evaluation index) was equated with the proportion of iterations with a higher score than our clustering solution.

#### Temporal modularity of functional networks

Following earlier work (Bassett et al. 2011), the maximum overlap discrete wavelet transform was used to decompose the time series for each ROI into wavelet coefficients in the range of 0.06 - 0.12Hz (Haar wavelet family, scale 2). The decomposed time series were divided into windows of *w*=64 seconds (32 imaging volumes; corresponding to 2 bins of 8 trials; see *Procedure* above) where contiguous windows overlapped by 1 bin, i.e. by 50% (Chai et al. 2016; Standage et al. 2020). We constructed functional networks in each time window by calculating the magnitude squared spectral coherence between each pair of brain regions and taking the mean coherence over the specified range of frequencies (Bassett et al. 2011). All coherence values less than the 95th percentile of a null model were set to zero. For each of baseline and learning, the null model was constructed over 10,000 iterations by selecting two regions and start times at random, shuffling their contents over window *w*, and calculating their coherence in the same way as for the original time series. This approach produced multislice networks with a mean sparsity of ∼80%. We determined the modular composition of the resulting multislice networks with a generalised Louvain method for time-resolved clustering (Jeub et al., 2019). This algorithm was repeated 100 times, resulting in 100 clustering solutions (a.k.a. partitions), each of which maximised a modularity ‘quality’ function (Mucha et al. 2010). On each repetition, we initialised the algorithm at random (uniform distribution) and iterated until a stable partition was found, *i*.*e*. the output on iteration *n* served as the input on iteration *n+1* until the output matched the input (Bassett et al. 2011; Mucha et al. 2010), where nodes were assigned to modules at random from all assignments that increased the quality function, and the probability of assignment was proportional to this increase. We used the standard spatial and temporal resolution parameters *γ* = 1 and *ω* = 1 respectively, and the standard null model by Newman and Girvan (Newman and Girvan 2004) for the quality function (Mucha et al. 2010; Sporns and Betzel 2016). Note that our results were qualitatively robust to variation of the above parameter values (see Supplementary Materials).

The flexibility of each region refers to the number of times it changes module allegiance, relative to the number of possible changes, i.e. flexibility = n_f_ / (T - 1), where n_f_ is the number of changes and T is the number of slices (time windows) in the multislice network (Bassett et al. 2011). We defined the flexibility for each partition as the mean flexibility of the regions, before calculating the mean across all partitions.

The cohesive flexibility (a.k.a. cohesion strength) of each region refers to the number of times it changes module allegiance together with each other region, summed over all other regions (Telesford et al. 2017). If *N*_*slice*_ is the number of time slices in a multislice network, *K(s)* and *M(s+1)* are the respective memberships of modules *k* and *m* at slices *s < N*_*slice*_ and *s+1*, and *G* is the set of regions moving from *k* to *m* at this time, then regions *i* and *j* move cohesively between modules *k* and *m* if and only if *{i,j}* ⊆ *K(s)* and {*i,j}* ⊆ *M(s+1)*. Defining such cohesive movements by *cs*_*i,j*_*(s, s+1) = 1 [0 otherwise]*, then cohesion strength is defined for each region *i* by *CS*_*i*_ *=* ∑_*s,s+1*_ ∑_*j ≠ i*_ *cs*_*i,j*_. We defined the cohesion strength for each partition as the mean cohesion strength of the regions, before calculating the mean across all partitions.

The disjointed flexibility (a.k.a. disjointedness) of each region refers to the number of times it changes modules independently, i.e. without other regions, relative to the number of possible changes. Following the above definitions, region *i* moves disjointedly if and only if *i* ∈ *K(s), I* ∈ *M(s+1)* and *G = {i}*. Defining such disjointed movements by *d*_*i*_*(s, s+1) = 1 [0 otherwise]* then disjointedness is defined for each region *i* by *D*_*i*_ *=* ∑_*s,s+1*_*d*_*i*_ */ (N*_*slice*_ *- 1)*. We defined the disjointedness for each partition as the mean disjointedness of the regions, before calculating the mean across all partitions.

#### Module allegiance matrix

We constructed a matrix *T*, where the elements *T*_*i,j*_ refer to the number of times regions *i* and *j* were assigned to the same module over all time slices, partitions and participants during a given task epoch. We then constructed the module allegiance matrix *P = (1/C)T*, where *C* is the total number of time slices in all of these partitions. Values that did not exceed those of a null model were set to 0 (Braun et al. 2015).

#### Clustering of the module allegiance matrix with non-negative matrix factorization

To derive summary networks from the module allegiance matrix, we used non-negative matrix factorisation [SymNMF; (Kuang, Ding, and Park 2012; Lee and Seung 1999)], which decomposes a symmetric matrix *Y* as *Y* = *AA*^*T*^, where *A* is a rectangular matrix containing a set of nonnegative factors that minimise a sum-of-squares error criterion. SymNMF has previously shown good performance in community detection problems (Kuang, Ding, and Park 2012; Kuang, Yun, and Park 2015), and has the benefit of producing a natural clustering solution by assigning each observation to the factor on which it loads most strongly. We used the Newton-like SymNMF algorithm described by Kuang et al. (2012), where for each number of factors between 2 and 15, we fit the model using 250 random initializations of the factor matrix H, with entries sampled from a uniform distribution on the interval [0, 1]. We then computed, for each rank, the average root mean squared reconstruction error, the dispersion coefficient (H. Kim and Park 2007), and the cophenetic correlation (Brunet et al. 2004). The latter two measures quantify the consistency of the clustering solutions returned by different initializations. We judged that a rank 5 solution was acceptable, as it explained over 80% of the variance in the observed data, it was acceptable by both the dispersion and cophenetic correlation criteria, and the generation of additional clusters provided little improvement in explained variance. We then selected the rank 5 solution with the lowest reconstruction error, and generated a clustering solution by assigning each brain region to the factor with the highest loading.

