## Supplementary material for "Whole-brain dynamics of human sensorimotor adaptation"

Dominic I. Standage*[1,3], Corson N. Areshenkoff[1], Daniel J. Gale[1], Joseph Y. Nashed[1,3], J. Randall Flanagan[1,2], Jason P. Gallivan*[1,2,3]

[1]Centre for Neuroscience Studies, [2]Department of Psychology, and [3]Department of Biomedical and Molecular Sciences, Queen's University, Kingston, Ontario, Canada

*Correspondence:

Dr. Dominic Standage
Department of Biomedical and Molecular Sciences, Centre for Neuroscience Studies
Queen's University


Dr. Jason Gallivan
Department of Psychology, and Department of Biomedical and Molecular Sciences
Queen's University


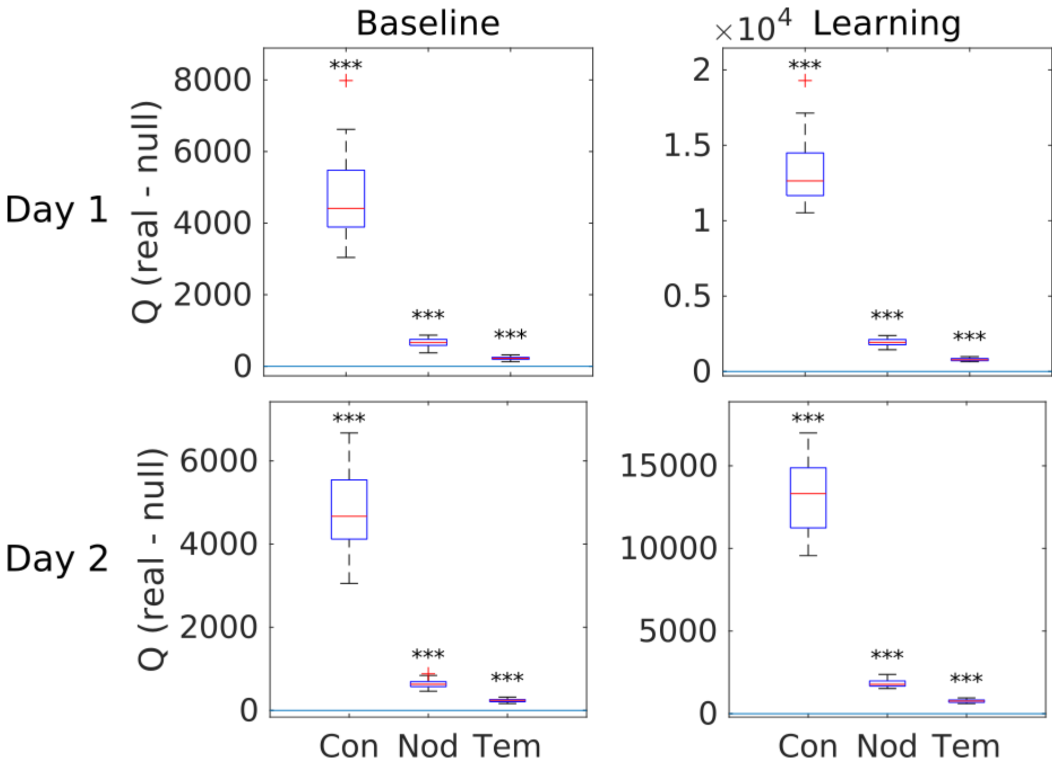


**Figure S1. Temporal networks showed significant modularity during baseline and learning on each day.** The difference in Q between real and permuted networks (real minus permuted) for connectional (Con), nodal (Nod) and temporal (Tem) null models (Mucha et al. 2010) was highly significant during baseline [top left, connectional: t(31) = 25.007, p = 4.104e-22; nodal: t(31) = 32.077, p = 2.417e-25; temporal: t(31) = 27.487, p = 2.475e-23] and learning [top right, connectional: t(31) = 36.672, p = 4.225e-27; nodal: t(31) = 45.654, p = 5.355e-30; temporal t(31) = 52.997, p = 5.752e-32] on Day 1 and during baseline [bottom left, connectional: t(31) = 29.079, p = 4.601e-24; nodal: t(31) = 38.350, p = 1.086e-27; temporal: t(31) = 33.480, p = 4.601e-24] and re-learning [bottom right, connectional: t(31) = 33.360, p = 7.408e-26; nodal: t(31) = 48.405, p = 8.941e-31; temporal t(31) = 46.180, p = 3.772e-30] on Day 2. As in the main text, three stars indicate p < 1e-3.

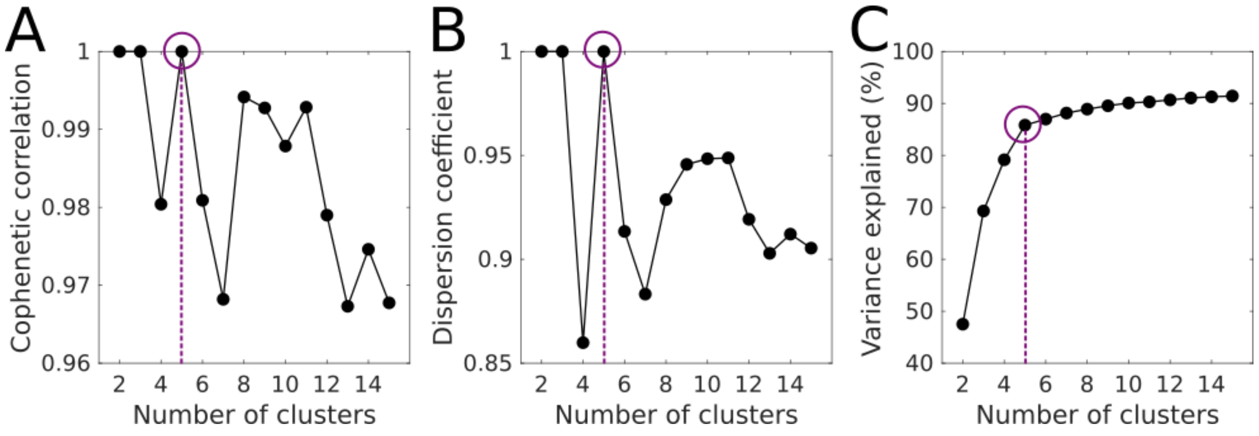

**Figure S2. Clustering diagnostics for symmetric non-negative matrix factorization of the module allegiance matrix.** Symmetric NMF was performed using 250 random initializations for factors 2 to 15. Regions were then assigned to a cluster corresponding to the factor with the greatest loading. For each number of factors, we computed the cophenetic correlation, dispersion, and explained variance. Dashed vertical line in each panel indicates the chosen solution of 5 clusters.

### Robustness of our results

Our results were robust to the width of the time windows used to construct temporal networks, and the proportion of learning defined as 'early' and 'late'. Below, we doubled the width of the time windows (for constructing temporal networks) from 64 seconds to 128 seconds, where early and late learning corresponded to windows 1 to 9 and 11 to 19 (out of 19 windows) respectively. Derivation of the summary networks and all statistical procedures for identifying networks of interest were identical to the main text. The similarity of these alternative summary networks to those in the main text was 100% (Network 1), 96% (Network 2), 88% (Network 3), 86% (Network 4) and 97% (Network 5), where similarity was defined for each pair of networks as the ratio of the size of their intersection to the size of their union.

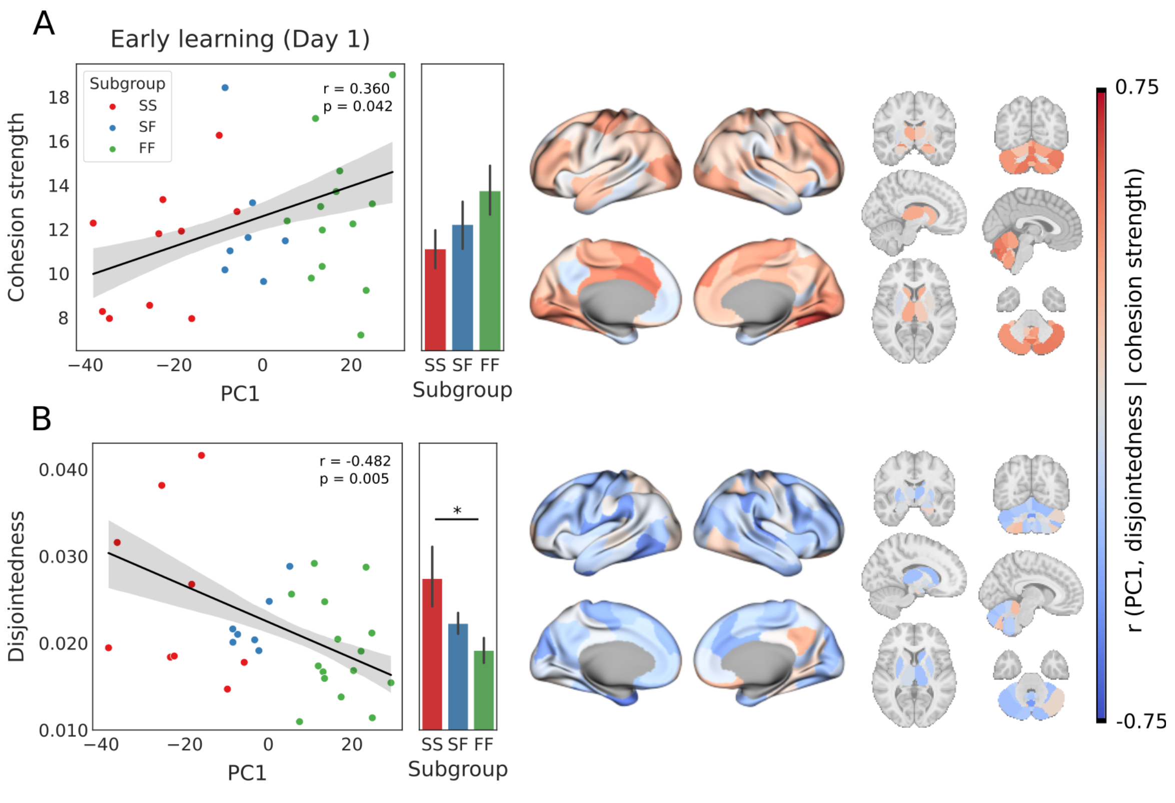


**Figure S3. Cohesion strength and disjointedness under alternative parameter values (cf. Figure 3, main text). (A)** Scatter plots show mean cohesion strength during early learning, plotted over PC1. Green, red and blue filled circles correspond to FF, SS and SF subgroups respectively. The fitted line shows the best linear fit, where the shaded area shows ±1 SE. Subgroup means ±1 SE are shown to the right of the scatter plots. Differences between the subgroups were non-significant [FF - SF: $t(20) = 0.853$, $p = 0.404$. FF - SS: $t(3) = 1.718$, $p = 0.099$. SF - SS: $t(15) = 0.781$, $p = 0.447$]. 126 out of 142 regions showed a positive correlation between PC1 and cohesion strength, 42 of which were significant ($0.351 < r < 0.669$, $3.0e\text{-}5 < p < 0.049$). None of the 16 negative correlations with PC1 were significant ($-0.147 < r <$

-5.801e-4, 0.420 < p < 0.998). **(B)** Mean disjointedness as a function of PC1 during early learning. SS was more disjointed than FF (FF > SS: t(20) = -2.373, p = 0.026), but SF did not differ significantly from SS [t(15) = -1.134, p = 0.275] or FF [t(20) = -1.311, p = 0.205]. Star in the bar plot indicates statistical significance (p < 0.05). 115 out of 142 regions showed a negative correlation between PC1 and disjointedness, 17 of which were significant (-0.6 < r < -0.357, 2.93e-4 < p < 0.046). None of the 27 negative correlations with PC1 were significant (0.006 > r > 0.315, 0.079 < p < 0.973). **(C)** Brain plots show correlations between PC1 and cohesion strength (upper plots) and disjointedness (lower plots) for each region. Cohesion strength and disjointedness are shown under a divergent colour scheme, ranging from strongly negative (dark blue) to strongly positive (dark red).

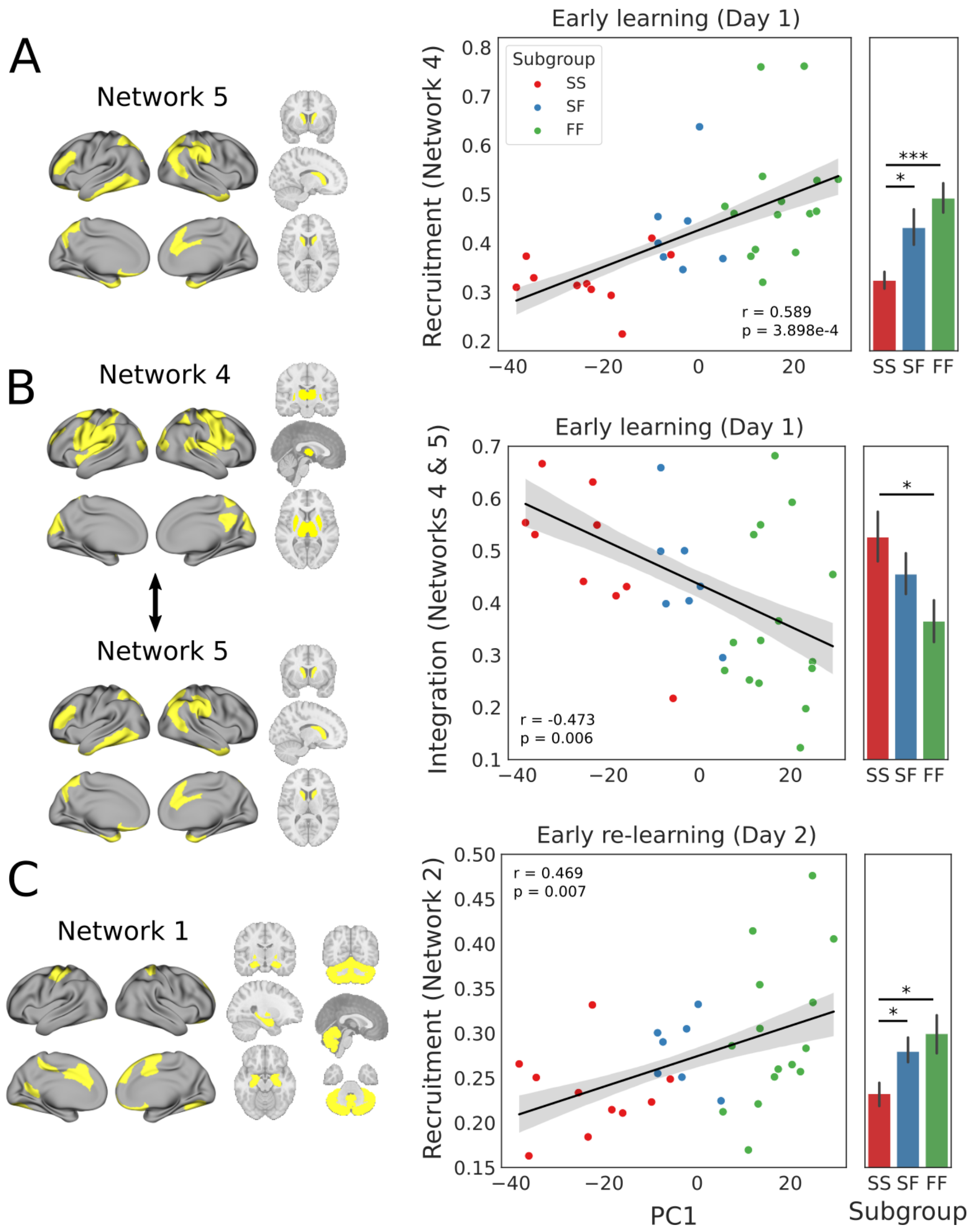

**Figure S4. Recruitment and integration under alternative parameter values (cf. Figure 5, main text). (A)** During early learning, recruitment of Network 5 was positively correlated with PC1, where recruitment by FF and SF participants was statistically indistinguishable [2-sample t-test, $t(20) = 1.112$, $p = 0.279$] but was greater among FF [$t(23) = 3.97$, $p = 6.061e-4$] and SF [$t(15) = 2.897$, $p = 0.011$] than SS (rightmost panel). **(B)** During early learning, integration between Networks 4 and 5 was negatively correlated with PC1 where integration between these networks was greater among FF participants than SS [$t(23) = -2.425$, $p = 0.024$] but did not differ statistically between FF and SF [$t(20) = -1.329$, $p = 0.199$] nor between SF and SS [$t(15) = -0.979$, $p = 0.343$] (rightmost panel). **(C)** During early re-learning, recruitment of Network 1 was positively correlated with PC1 where recruitment by FF and SF participants was statistically indistinguishable [$t(20) = 0.595$, $p = 0.558$] but was greater among FF [$t(23) = 2.299$, $p = 0.031$] and SF [$t(15) = 2.217$, $p = 0.043$] participants than SS (rightmost panel).. In scatter plots, fitted line shows linear fit, where shading corresponds to ±1 SE. Bar plots show means, where error bars show ±1 SE. In bar plots, stars indicate statistical significance (one star: $p < 0.05$; three stars: $p < 1e-3$).

#### The association between faster learning and recruitment of Network 1 remained the case when summary networks were derived during early re-learning

Summary networks in the main text were derived during early learning (Day 1). Because recruitment of Network 1 during early *re*-learning (Day 2) was associated with a faster learning profile, we sought to determine whether this result would remain the case if we derived the summary networks during early re-learning, i.e. during the same epoch as we calculated recruitment on Day 2. We therefore derived summary networks during early re-learning (64 second windows, as in the main text). We then calculating the correlation between PC1 and recruitment of each network during early re-learning, correcting for multiple comparisons (same procedure as in the main text). The new ‘Network 1’ was 88% similar to Network 1 in the main text and was the only network in which recruitment during early re-learning was significantly correlated with PC1. Like the original Network 1, this network included bilateral hippocampus, striatum and the entire cerebellum. Furthermore, recruitment of this network by the FF and SF subgroups was statistically indistinguishable, and was greater in both of these subgroups than in the SS subgroup. Thus, our Day 2 results did not rely on deriving the summary networks on Day 1.

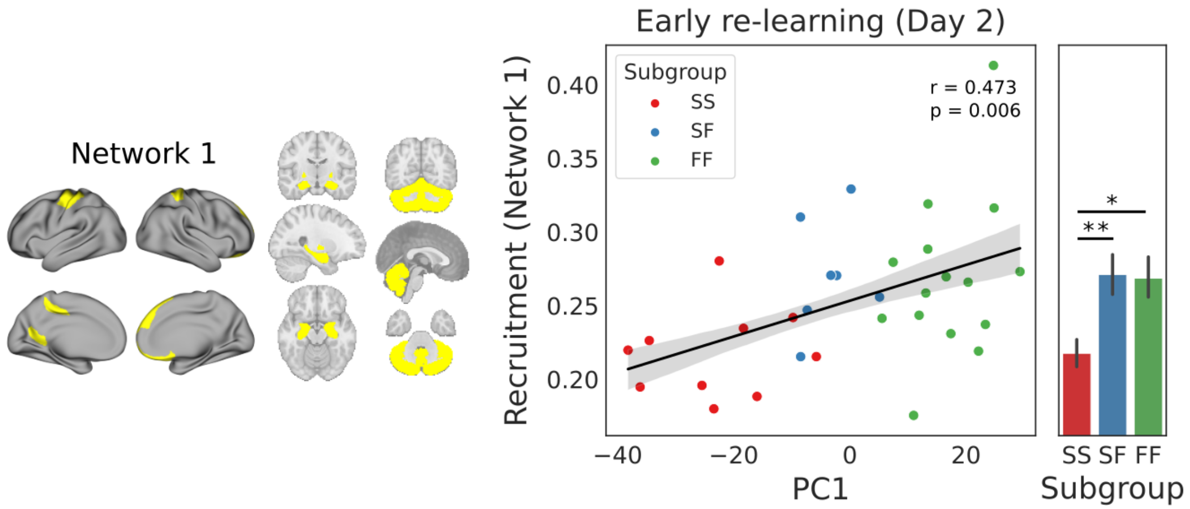


**Figure S5. Greater recruitment of Network 1 during early re-learning was associated with a faster learning profile, whether the summary networks were derived during early learning (main text) or during early re-learning (cf. Figure 5C, main text).** During early re-learning, recruitment of 'Network 1' (derived during early re-learning and composed mainly of hippocampal, striatal and cerebellar regions, left side) was positively correlated with PC1, where recruitment by FF and SF participants was statistically indistinguishable [t(20) = 0.108, p = 0.915] but was greater among FF [t(23) = 2.697, p = 0.013] and SF [t(15) = 3.226, p = 0.006 participants than SS (rightmost panel). In scatter plots, fitted line shows linear fit, where shading corresponds to ±1 SE. Bar plots show means, where error bars show ±1 SE. In bar plots, stars indicate statistical significance (one star: p < 0.05; two stars: p < 0.01).

Finally, having determined that our results for re-learning (Figure 5C, main text; Figure S5 above) did not depend on whether we derived the summary networks during early learning or re-learning, we re-ran our analysis of recruitment and integration during early learning, using the summary networks derived during early re-learning above. Results of all statistical tests were the same as in the main text. There was a significant positive correlation between PC1 and the recruitment of 'Network 5' (82% similar to Network 5 in the main text) during early learning (r = 0.493, p = 0.004, adjusted p = 0.021). This network was the only one whose recruitment was significantly correlated with PC1, and its recruitment by FF and SF participants was statistically indistinguishable [2-sample t-test, t(20) = 0.034, p = 0.973] but was greater among FF [t(23) = 3.646, p = 0.001] and SF [t(15) = 4.616, p = 3.336e-4] than SS participants. Integration of 'Network 4' (83% similar to Network 4 in the main text) and 'Network 5' was negatively correlated with PC1 (r = -0.485, p = 0.005, adjusted p = 0.020] where integration between these networks was greater among FF participants than SS [t(23) = -2.290, p = 0.032] but did not differ statistically between FF and SF [t(20) = -0.378, p = 0.709] nor between SF and SS [t(20) = -1.930, p = 0.073]. As in the main text, no other pair of networks showed a significant correlation between their integration and PC1 during early learning. Thus, all of our results based on the summary networks in the main text (derived during early learning) remained the case when we derived the summary networks during early re-learning.

#### Table S1. Summary network composition (networks in main text)

**Network 1**

| *Region* | *Atlas* |
|---|---|
| 7Networks_LH_Vis_2 | Schaefer |
| 7Networks_LH_SomMot_5 | Schaefer |
| 7Networks_LH_SalVentAttn_Med_1 | Schaefer |
| 7Networks_LH_SalVentAttn_Med_2 | Schaefer |
| 7Networks_LH_Default_pCunPCC_1 | Schaefer |
| 7Networks_RH_Vis_2 | Schaefer |
| 7Networks_RH_SomMot_7 | Schaefer |
| 7Networks_RH_SalVentAttn_Med_2 | Schaefer |
| 7Networks_RH_Limbic_OFC_1 | Schaefer |
| 7Networks_RH_Default_PFCdPFCm_2 | Schaefer |
| 7Networks_RH_Default_PFCdPFCm_3 | Schaefer |
| L-Pallidum | Harvard-Oxford sub-cortical |
| L-Hippocampus | Harvard-Oxford sub-cortical |
| L-Amygdala | Harvard-Oxford sub-cortical |
| L-Accumbens | Harvard-Oxford sub-cortical |
| R-Pallidum | Harvard-Oxford sub-cortical |
| R-Hippocampus | Harvard-Oxford sub-cortical |
| R-Amygdala | Harvard-Oxford sub-cortical |
| R-Accumbens | Harvard-Oxford sub-cortical |
| L-I-IV | Diedrichsen (SUIT) cerebellum |
| R-I-IV | Diedrichsen (SUIT) cerebellum |
| L-V | Diedrichsen (SUIT) cerebellum |
| R-V | Diedrichsen (SUIT) cerebellum |
| L-VI | Diedrichsen (SUIT) cerebellum |
| VI | Diedrichsen (SUIT) cerebellum |
| R-VI | Diedrichsen (SUIT) cerebellum |
| L-Crus | Diedrichsen (SUIT) cerebellum |
| Crus | Diedrichsen (SUIT) cerebellum |
| R-Crus | Diedrichsen (SUIT) cerebellum |
| L-Crus | Diedrichsen (SUIT) cerebellum |
| Crus | Diedrichsen (SUIT) cerebellum |
| R-Crus | Diedrichsen (SUIT) cerebellum |
| L-VIIb | Diedrichsen (SUIT) cerebellum |
| VIIb | Diedrichsen (SUIT) cerebellum |
| R-VIIb | Diedrichsen (SUIT) cerebellum |
| L-VIIIa | Diedrichsen (SUIT) cerebellum |
| VIIIa | Diedrichsen (SUIT) cerebellum |
| R-VIIIa | Diedrichsen (SUIT) cerebellum |
| L-VIIIb | Diedrichsen (SUIT) cerebellum |
| VIIIb | Diedrichsen (SUIT) cerebellum |
| R-VIIIb | Diedrichsen (SUIT) cerebellum |
| L-IX | Diedrichsen (SUIT) cerebellum |
| IX | Diedrichsen (SUIT) cerebellum |
| R-IX | Diedrichsen (SUIT) cerebellum |
| L-X | Diedrichsen (SUIT) cerebellum |
| X | Diedrichsen (SUIT) cerebellum |
| R-X | Diedrichsen (SUIT) cerebellum |

**Network 2**

| *Region* | *Atlas* |
|---|---|
| 7Networks_LH_SalVentAttn_FrOperIns_2 | Schaefer |
| 7Networks_LH_SalVentAttn_Med_3 | Schaefer |
| 7Networks_LH_Cont_Par_1 | Schaefer |
| 7Networks_LH_Cont_Cing_1 | Schaefer |
| 7Networks_LH_Default_Temp_1 | Schaefer |
| 7Networks_LH_Default_Temp_2 | Schaefer |
| 7Networks_LH_Default_Par_1 | Schaefer |
| 7Networks_LH_Default_Par_2 | Schaefer |
| 7Networks_LH_Default_PFC_1 | Schaefer |
| 7Networks_LH_Default_PFC_2 | Schaefer |
| 7Networks_LH_Default_PFC_3 | Schaefer |
| 7Networks_LH_Default_PFC_4 | Schaefer |
| 7Networks_LH_Default_PFC_5 | Schaefer |
| 7Networks_LH_Default_PFC_6 | Schaefer |
| 7Networks_LH_Default_pCunPCC_2 | Schaefer |
| 7Networks_RH_SalVentAttn_Med_1 | Schaefer |
| 7Networks_RH_Cont_PFCl_1 | Schaefer |
| 7Networks_RH_Cont_PFCl_2 | Schaefer |
| 7Networks_RH_Cont_PFCl_4 | Schaefer |
| 7Networks_RH_Cont_Cing_1 | Schaefer |
| 7Networks_RH_Cont_pCun_1 | Schaefer |
| 7Networks_RH_Default_Par_1 | Schaefer |
| 7Networks_RH_Default_Temp_1 | Schaefer |
| 7Networks_RH_Default_Temp_2 | Schaefer |
| 7Networks_RH_Default_Temp_3 | Schaefer |
| 7Networks_RH_Default_PFCv_1 | Schaefer |
| 7Networks_RH_Default_PFCv_2 | Schaefer |
| 7Networks_RH_Default_pCunPCC_1 | Schaefer |

**Network 3**

| *Region* | *Atlas* |
|---|---|
| 7Networks_LH_Vis_1 | Schaefer |
| 7Networks_LH_Vis_3 | Schaefer |
| 7Networks_LH_Vis_4 | Schaefer |
| 7Networks_LH_Vis_5 | Schaefer |
| 7Networks_LH_Vis_6 | Schaefer |
| 7Networks_LH_Vis_7 | Schaefer |
| 7Networks_LH_SomMot_6 | Schaefer |
| 7Networks_LH_Default_PFC_7 | Schaefer |
| 7Networks_RH_Vis_1 | Schaefer |
| 7Networks_RH_Vis_3 | Schaefer |
| 7Networks_RH_Vis_4 | Schaefer |
| 7Networks_RH_Vis_5 | Schaefer |
| 7Networks_RH_Vis_6 | Schaefer |
| 7Networks_RH_SomMot_8 | Schaefer |
| 7Networks_RH_Default_PFCdPFCm_1 | Schaefer |

**Network 4**

| *Region* | *Atlas* |
|---|---|
| 7Networks_LH_Vis_8 | Schaefer |
| 7Networks_LH_Vis_9 | Schaefer |
| 7Networks_LH_SomMot_1 | Schaefer |
| 7Networks_LH_SomMot_2 | Schaefer |
| 7Networks_LH_SomMot_3 | Schaefer |
| 7Networks_LH_SomMot_4 | Schaefer |
| 7Networks_LH_DorsAttn_Post_4 | Schaefer |
| 7Networks_LH_DorsAttn_Post_6 | Schaefer |
| 7Networks_LH_DorsAttn_PrCv_1 | Schaefer |
| 7Networks_LH_DorsAttn_FEF_1 | Schaefer |
| 7Networks_LH_SalVentAttn_ParOper_1 | Schaefer |
| 7Networks_LH_SalVentAttn_FrOperIns_1 | Schaefer |
| 7Networks_LH_SalVentAttn_PFCl_1 | Schaefer |
| 7Networks_RH_Vis_7 | Schaefer |
| 7Networks_RH_Vis_8 | Schaefer |
| 7Networks_RH_SomMot_1 | Schaefer |
| 7Networks_RH_SomMot_2 | Schaefer |
| 7Networks_RH_SomMot_3 | Schaefer |
| 7Networks_RH_SomMot_4 | Schaefer |
| 7Networks_RH_SomMot_5 | Schaefer |
| 7Networks_RH_SomMot_6 | Schaefer |
| 7Networks_RH_DorsAttn_Post_3 | Schaefer |
| 7Networks_RH_DorsAttn_Post_5 | Schaefer |
| 7Networks_RH_DorsAttn_PrCv_1 | Schaefer |
| 7Networks_RH_DorsAttn_FEF_1 | Schaefer |
| 7Networks_RH_SalVentAttn_TempOccPar_1 | Schaefer |
| 7Networks_RH_Default_pCunPCC_2 | Schaefer |
| L-Thalamus | Harvard-Oxford sub-cortical |
| L-Putamen | Harvard-Oxford sub-cortical |
| R-Thalamus | Harvard-Oxford sub-cortical |
| R-Putamen | Harvard-Oxford sub-cortical |

**Network 5**

| *Region* | *Atlas* |
|---|---|
| 7Networks_LH_DorsAttn_Post_1 | Schaefer |
| 7Networks_LH_DorsAttn_Post_2 | Schaefer |
| 7Networks_LH_DorsAttn_Post_3 | Schaefer |
| 7Networks_LH_DorsAttn_Post_5 | Schaefer |
| 7Networks_LH_Limbic_OFC_1 | Schaefer |
| 7Networks_LH_Limbic_TempPole_1 | Schaefer |
| 7Networks_LH_Limbic_TempPole_2 | Schaefer |
| 7Networks_LH_Cont_PFCl_1 | Schaefer |
| 7Networks_LH_Cont_pCun_1 | Schaefer |
| 7Networks_RH_DorsAttn_Post_1 | Schaefer |
| 7Networks_RH_DorsAttn_Post_2 | Schaefer |
| 7Networks_RH_DorsAttn_Post_4 | Schaefer |
| 7Networks_RH_SalVentAttn_TempOccPar_2 | Schaefer |
| 7Networks_RH_SalVentAttn_FrOperIns_1 | Schaefer |
| 7Networks_RH_Limbic_TempPole_1 | Schaefer |
| 7Networks_RH_Cont_Par_1 | Schaefer |
| 7Networks_RH_Cont_Par_2 | Schaefer |
| 7Networks_RH_Cont_PFCl_3 | Schaefer |
| 7Networks_RH_Cont_PFCmp_1 | Schaefer |
| L-Caudate | Harvard-Oxford sub-cortical |
| R-Caudate | Harvard-Oxford sub-cortical |
